# Quantitative Prevalence, Chemical Speciation, and ECM-Embedded Networks of Biogenic Silicon in Animal Tissues Revealed by Refined Analytical Methods

**DOI:** 10.1101/2025.10.02.679922

**Authors:** Yanjun Wang, Zhiyao Lu, Ruoyu Yao, Guangxin Zhou, Wenlu Meng, Yulian Wang, Chunyan Liu, Siyu Li, Zhuofan Wu, Qingyun Wang, Zhen Xing, Chunming Wang, Yuan Yin, Rui Wang, Lei Dong

**Author notes:** Corresponding author. (L.D.); (R.W.); (Y.Y.).

## Abstract

Silicon, abundant in Earth’s biosphere, plays crucial roles in many organisms. However, its biological functions in animals remain inadequately characterized, primarily constrained by methodological limitations in quantifying, identifying, and visualizing biogenic silicon in tissues. Here, we refined the molybdenum blue colorimetric method (MB) for more accurate silicon quantification, developed integrated mass spectrometry techniques to delineate chemical natures of biogenic silicon, and optimized micro-X-ray fluorescence (micro-XRF) imaging for spatial mapping in animals. Employing these methodologies, we demonstrated that silicon abundance in animals substantially exceeds prior estimates—surpassing essential elements like iron. We also revealed the chemical architectures of diverse silicon compounds and visualized tissue-specific distributions in mice and humans, indicating its predominant localization within extracellular matrix (ECM) and suggesting the formation of silicified networks. These findings establish silicon as a structurally and quantitatively significant element with profound biological implications, warranting its recognition as an essential factor in human physiology and pathology.

## Introduction

Silicon ranks among the most abundant elements in Earth’s biosphere. Silicon compounds in the Earth’s crust and soil play fundamental roles in the origin and evolution of life^1–3^. Recent studies suggest that silicates may catalyze the synthesis of critical biomolecules such as amino acids and carbohydrates. This catalytic capacity of silicon compounds likely proved pivotal in the emergence of early life forms and continues to exert biological influence in contemporary terrestrial organisms^4–6^. The biological significance of silicon compounds manifests across diverse organisms. For instance, marine diatoms and sponges incorporate silica into their structural frameworks^7,8^, while plants actively assimilate silicon from soil, accumulating it as phytoliths comprising 0.1%-10% of dry weight^9,10^. This silicon accumulation enhances plant resilience against environmental stressors and pathogens, demonstrating silicon’s regulatory capacity in complex biological systems^11–13^.

In animals, particularly mammals, silicon’s biological roles have gained increasing attention yet remain poorly understood^14–16^. The seminal 1972 study by Carlisle and Schwarz revealed that silicon deficiency induces severe developmental impairments in chicks and rats, establishing silicon’s essentiality in animal development^17,18^. Contemporary research implicates silicon in physiological and pathological processes including collagen synthesis, bone mineralization, and neurodegenerative disease modulation^19–26^. However, despite these critical findings, systematic investigations into the precise biological regulatory mechanisms of silicon compounds in animals remain conspicuously lacking.

The principal challenges in studying silicon compounds in animal tissues stem from current analytical limitations. First, while inductively coupled plasma optical emission spectrometry (ICP-OES) offers high sensitivity for silicon quantification, its application to biological samples is constrained by the inherent incompatibility of silicon compounds with the acidic pretreatment conditions required for analysis, a limitation that may systematically underestimate tissue silicon content and obscure its biological relevance^18,21^. The molybdenum blue (MB) colorimetric method, successfully employed in plant studies, provides a potential solution for animal tissue silicon quantification but requires methodological adaptation^7,12,15,27–32^.

Second, elucidating the chemical speciation of silicon in biological systems presents another critical challenge. While silicon predominantly exists as silica in marine organisms and as diversified phytoliths in plants^8,9,33^, its chemical forms in animal tissues remain enigmatic. This knowledge gap arises from both the ICP-OES-based underestimation of tissue silicon content and the analytical difficulties inherent in acid-dependent extraction protocols. Traditional assumptions of monomeric silicate predominance in animals likely misrepresent reality, given silicon’s propensity for polymerization and potential interactions with biomolecules in complex physiological environments^17,18^. Recent advancements in high-sensitivity mass spectrometry provide unprecedented opportunities to characterize silicon species in animal tissues^34,35,44–46^.

Finally, functional analysis of specific molecules in animal tissues typically relies on visualization techniques requiring specific molecular markers - an approach fundamentally impeded for silicon compounds. Unlike most biomolecules, silicon compounds exhibit chemical inertness and low immunogenicity, rendering conventional histochemical staining and immunolabeling techniques ineffective for silicon imaging. Breakthroughs in high-resolution multi-element X-ray spectroscopy offer promising alternatives for silicon visualization in biological tissues^36–42^.

This study addresses these challenges by developing three complementary analytical approaches: (1) optimized MB method for accurate silicon quantification in animal tissues; (2) advanced mass spectrometry for chemical speciation analysis of tissue silicon; (3) micro-X-ray fluorescence (micro-XRF) imaging for spatial distribution mapping of silicon compounds. Through integrating these methodological innovations with systematic tissue analysis, we have clarified silicon’s authentic abundance, chemical nature, and distribution patterns in animal tissues. These findings advance our understanding of silicon’s biological functions and provide valuable insights to guide future investigations into silicon compound-mediated pathophysiological mechanisms and therapeutic applications.

## Results

### Modified HF-MB Method Reveals High Silicon Levels in Animal Tissues

Inductively coupled plasma optical emission spectrometry (ICP-OES) is commonly used for silicon quantification in animal tissues. However, the strong acid digestion process employed in sample pretreatment leads to silicon precipitation and measurement inaccuracies. Alkali and hydrofluoric acid, which can effectively dissolve silicon compounds, are prohibited from use due to their corrosive effects on the glass components of ICP-OES equipment. This indicates that the silicon content measured by ICP-OES in animal tissues may substantially deviate from true values. To address this issue, we modified the molybdenum blue (HF-MB) colorimetric method for silicon quantification by incorporating a hydrofluoric acid dissolution step (Fig. 1a) to ensure complete solubilization of silicon compounds. As shown in Fig. 1b, visible precipitates were observed in silicon standards (polysilicic acid and sodium silicate) and mouse tissues (liver and bone) processed with conventional acid digestion, whereas the HF-MB protocol yielded clear solutions after tissue digestion. Validation using gradient sodium silicate solutions with known levels revealed significant deviations from true values and severe nonlinearity in ICP-OES measurements. In contrast, the HF-MB method accurately quantified silicon levels in gradient sodium silicate solutions, with deviations below ±0.3 μg and excellent linearity (Fig. 1c).

**Figure 1.**
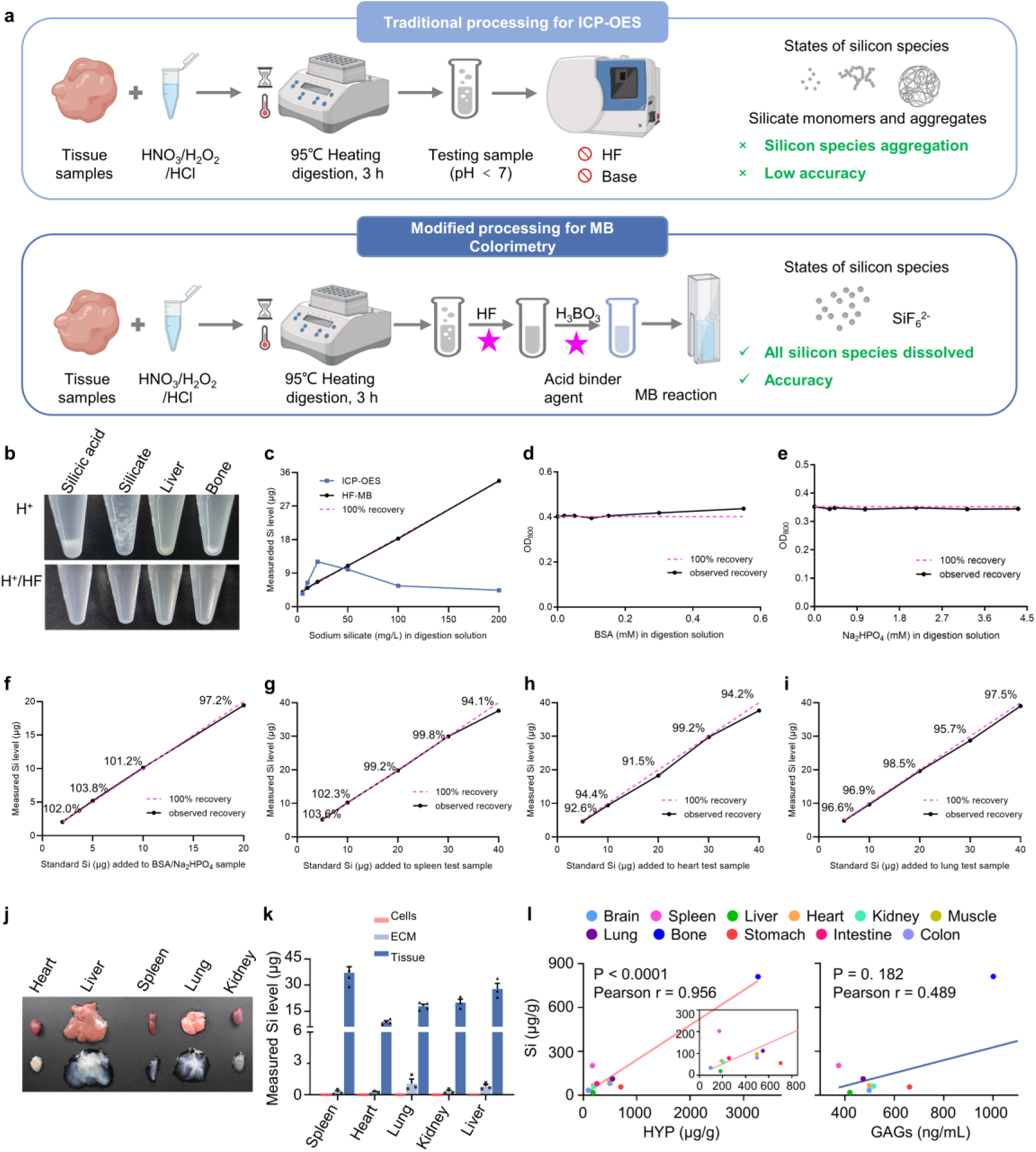
Quantification and analysis of silicon concentrations in mouse and human tissues. **a**, Schematic diagram of the silicon contents detection methods in animal tissues. **b**, Silicon aggregation in polysilicic acid, sodium silicate, mouse liver, and bone during various processing methods, which are the conventional processing for ICP-OES (acid digestion, HNO_3_/ H_2_O_2_/ HCl, 95°C, 3 h) and modified processing for HF-MB (dissolution of silicon species by hydrofluoric acid (HF) after acid digestion). **c**, Comparison of silicon levels in sodium silicate solutions after different processing determined by HF-MB and ICP-OES methods. **d**, **e**, Anti-interference test for silicate (5 μg) in 150 μL digestion solution measured by HF-MB. **f**-**i**, Spike-recovery experiments: addition of standard Si solutions to albumin/phosphate mixtures (BSA 0.55 mM, Na_2_HPO_4_ 4.31 mM, 150 μL digestion solution), or mouse tissues (spleen, heart, lung) (3 mg dry tissue, 150 μL digest solution), measured by HF-MB Method. **j**, Entire decellularized ECM of mouse tissues. **k**, Silicon levels in cells and ECM of mouse tissues (dry weight) determined by HF-MB. **l**, Relationship between silicon (determined by HF-MB) and hydroxyproline (HYP) or glycosaminoglycans (GAGs) concentrations in mouse tissues, with regression lines and Pearson correlation coefficients (r) and P values shown.

Subsequently, we analyzed potential interference from proteins and phosphorus in animal tissues on the HF-MB assay. Experimental results demonstrated that neither bovine serum albumin (0-0.55 mM, 150 μL digestion solution) nor disodium hydrogen phosphate (0-4.31 mM, 150 μL digestion solution) affected measurement accuracy (Fig. 1d and Fig. 1e). Additionally, spike-recovery experiments conducted in albumin/phosphate mixtures and mouse tissues (spleen, heart, and lung) demonstrated that the HF-MB method maintained consistent silicon extraction efficiency from complex biological matrices, with recovery rates consistently falling within the 90%-110% range (Figs. 1f-1i).

Following confirmation of the HF-MB method’s accuracy and reliability, we comprehensively compared tissue silicon levels in mice using both detection approaches (Table 1). ICP-OES measurements yielded silicon concentrations of 0.4-25 μg Si/g (wet weight), consistent with literature reports but showing substantial discrepancies from HF-MB results: HF-MB detected silicon concentrations exceeding ICP-OES values by over 10-fold in almost all tissues, more than 30-fold in brain, liver, and kidney, and an extraordinary 100-fold difference in muscle and bone. HF-MB measurements revealed silicon concentrations of 100-900 μg Si/g (wet weight) in bone, bone marrow, spleen, and lung, and 10-100 μg Si/g (wet weight) in other tissues, approaching the upper limit of conventionally defined “trace element” levels in animals.

**Table 1.** Quantification of silicon in mouse and human tissues and serum determined by HF-MB and ICP-OES method (n = 3 unless otherwise noted).

| Mouse tissues | ICP-OES (µg/g, wet weight) | ICP-OES (µg/g, dry weight) | HF-MB (µg/g, wet weight) | HF-MB (µg/g, dry weight) | Mouse tissues | ICP-OES (µg/g, wet weight) | ICP-OES (µg/g, dry weight) | HF-MB (µg/g, wet weight) | HF-MB (µg/g, dry weight) | Human tissues | ICP-OES (µg/g, wet weight) | ICP-OES (µg/g, dry weight) | HF-MB (µg/g, wet weight) | HF-MB (µg/g, dry weight) |
| --- | --- | --- | --- | --- | --- | --- | --- | --- | --- | --- | --- | --- | --- | --- |
| Spleen | 9.4±5.7 | 46.3±28.1 | 203.1±47.6 | 995.2±233.2 | Lung | 6.7±2.7 | 33.8±13.4 | 112.4±15.0 | 562.2±75.2 | Muscle | 7.4±2.1 | 36.0±10.9 | 107.5±8.7 | 506.4±19.0 |
| Brain | 1.0±0.2 | 5.4±1.3 | 34.3±1.4 | 185.3±7.4 | Colon | 7.6±3.1 | 46.2±19.0 | 80.6±24.6 | 491.5±150.1 | Liver | 14.8±5.7 | 49.4±19.0 | 179.1±68.0 | 586.4±169.1 |
| Muscle | 1.5±0.5 | 8.0±2.7 | 97.0±12.2 | 504.6±63.6 | Kidney | 2.5±0.5 | 11.2±2.4 | 61.7±10.4 | 277.6±46.9 | Stomach (n=1) | 12.3 | 43.9 | 221.8 | 793.1 |
| Liver | 0.4±0.03 | 1.5±0.1 | 19.2±3.8 | 71.0±14.1 | Intestine | 5.5±1.0 | 34.9±6.4 | 78.9±26.9 | 504.8±172.3 | Colon (n=4) | 12.3±1.7 | 68.9±9.8 | 151.2±19.6 | 846.3±109.6 |
| Pancreas | 3.6±2.0 | 15.3±8.7 | 24.3±10.6 | 104.2±45.5 | Bone | 4.7±1.3 | 8.9±2.6 | 811.2±128.8 | 1541.3±244.8 | Intestine | 6.9±1.1 | 44.4±3.1 | 62.5±23.9 | 400.0±132.5 |
| Stomach | 4.4±2.0 | 21.1±8.4 | 56.4±5.4 | 242.4±23.0 | Bone marrow | 24.7±12.9 | 236.4±186.1 | 245.6±56.7 | 1860.9±606.6 | Bone | 8.9±1.7 | 17.3±2.2 | 414.8±52.4 | 803.8±102.1 |
| Heart | 4.7±0.8 | 20.6±3.4 | 66.1±12.2 | 290.7±53.7 | Serum | 3.3±0.4 (µg/mL) |  | 64.1±7.9 (µg/mL) |  | Serum | 4.0±1.1 (µg/mL) |  | 54.4±2.3 (µg/mL) |  |

Similar discrepancies between the two methods were observed in human tissue analyses (Table 1). Notably, species-specific differences in silicon content were identified: human liver, stomach, and colon tissues contained substantially higher silicon levels than corresponding mouse tissues, whereas human bone showed lower silicon content than mouse bone. HF-MB analysis of common food sources revealed that meat products, particularly beef and lamb, contained higher silicon concentrations than vegetables (Table S1).

To investigate silicon distribution between extracellular and intracellular compartments, we employed whole-organ decellularization to obtain extracellular matrix (ECM) from mouse heart, liver, spleen, lung, and kidney (Fig. 1j), followed by collagenase digestion to isolate cellular components. HF-MB analysis demonstrated negligible intracellular silicon levels, with detectable silicon predominantly localized in the ECM, despite substantial silicon loss during processing (Fig. 1k). The observed silicon leakage during decellularization suggests that silicon associates with ECM components through non-covalent interactions rather than strong binding. Correlation analysis between silicon content and ECM components revealed a strong positive association with collagen content (as indicated by hydroxyproline levels; Pearson r = 0.956; P < 0.0001) but no correlation with glycosaminoglycans (GAGs) across multiple tissues (Fig. 1l).

These findings demonstrate that conventional ICP-OES severely underestimates silicon content in animal tissues, with actual levels exceeding previous reports by 1-2 orders of magnitude. The measured silicon abundance approaches or even surpasses that of iron (the most abundant trace element in humans; Fig. S1). The unexpectedly high tissue concentrations and specific ECM localization of silicon suggest potentially critical, yet previously unrecognized, biological functions.

### Diverse Chemical Nature of Biogenic Silicon in Animal Tissues Uncovered by Mass Spectrum

To investigate the chemical structures of biogenic silicon in animals, we employed mass spectrometry, following the strategy outlined in Fig. 2a. We first analyzed the proportions of soluble and insoluble silicon. By centrifuging tissue homogenates, we separated these two forms in mouse tissues and quantified them using HF-MB analysis. Soluble silicon consistently exceeded insoluble silicon across almost all tissues, with a relatively constant ratio between the two forms (Fig. 2b).

**Figure 2.**
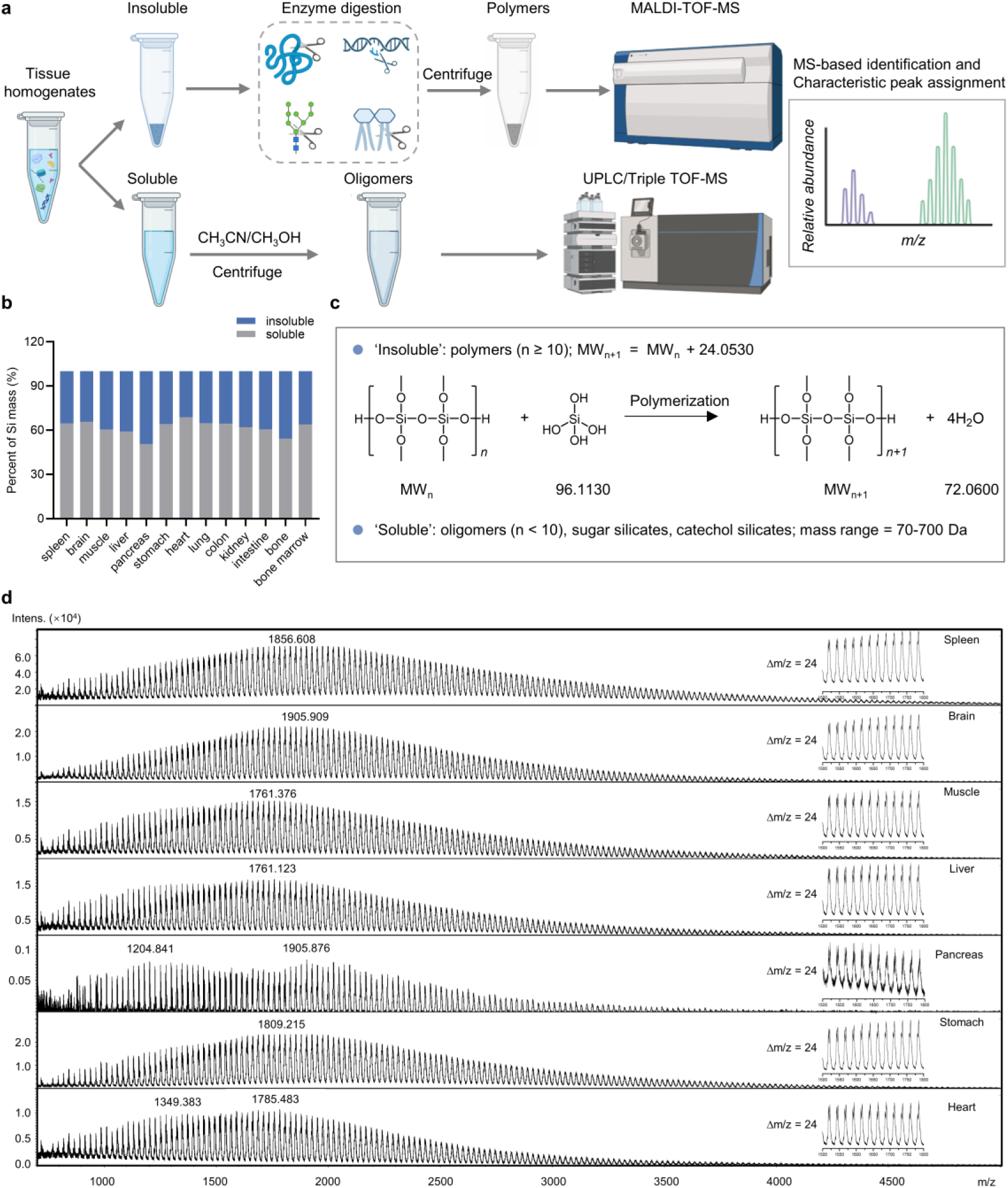
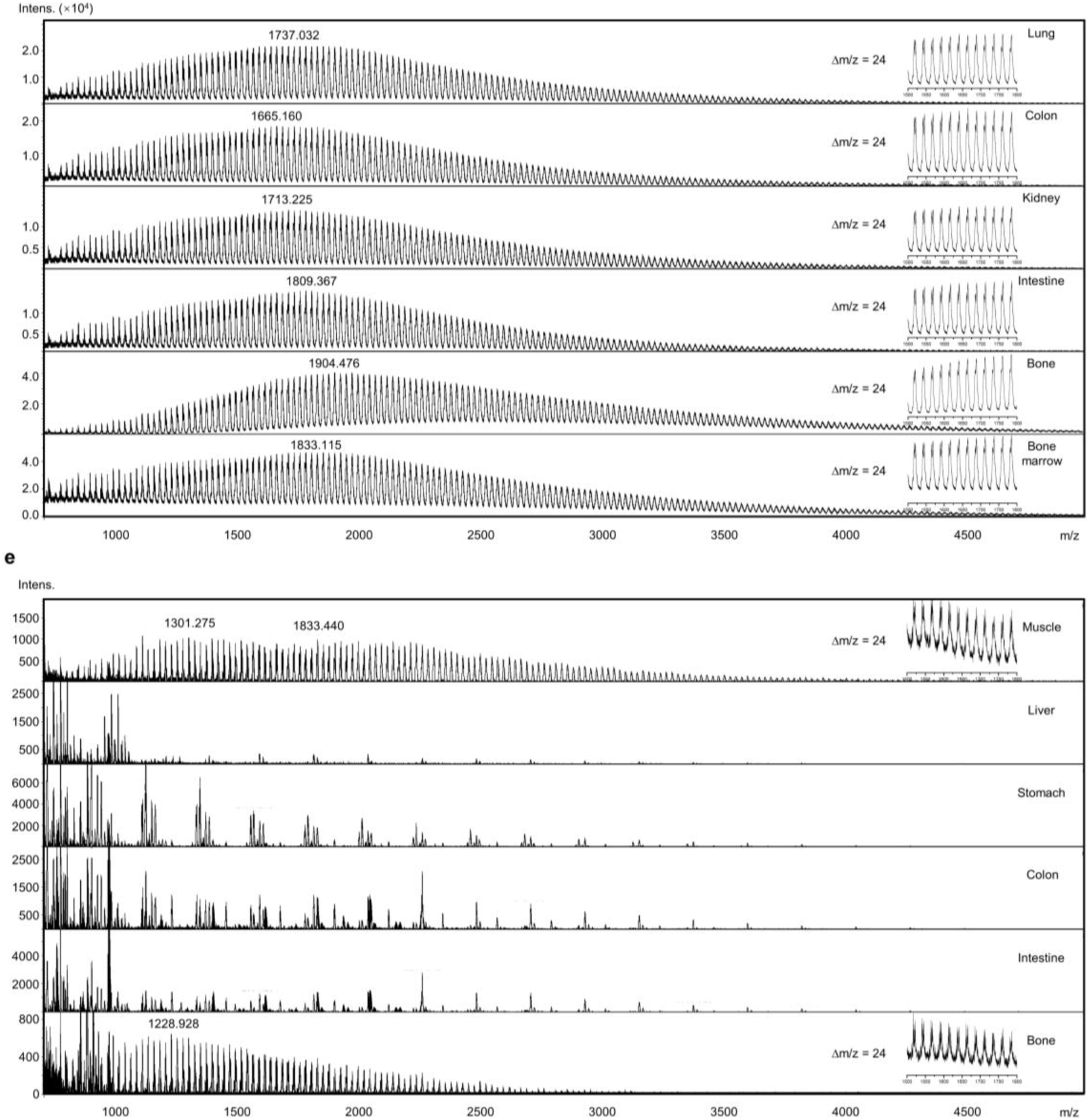
Identification of the chemical nature of biogenic silicon in mouse and human tissues. **a**, Schematic of the mass spectrum experimental setup for analyzing the chemical nature of biogenic silicon in tissue samples. The tissue homogenates were separated into the insoluble and soluble matter for analysis. It was hypothesized that soluble silicate oligomers are present in the supernatant, which were detected using UPLC/Triple-TOF-MS. 2 mM soluble sodium silicate solution was used as the synthetic standard sample for the oligomers. Conversely, insoluble silicate polymers were found in the precipitate. To enrich for silicate polymers, enzymatic digestion was first performed using protease, glycosidase, DNase, and lipase to remove biological macromolecules (proteins, polysaccharides, nucleic acids, and lipids), followed by washing away inorganic salts with water. MALDI-TOF-MS was then employed to detect the silicate polymers, with 1% (w/w) silicate polymers used as the synthetic standard sample. All synthetic silicate samples were prepared by adding hydrochloric acid to sodium silicate at the appropriate concentration, adjusting the pH to 7.4. **b**, Percent of Si mass in the insoluble and soluble matter of mouse tissue homogenates. **c**, Potential chemical nature of ‘insoluble’ and ‘soluble’ silicon species in animal tissues, MW: molecular weight. n: silicon number. **d**, Potential silicate polymers in mouse tissue samples identified by MALDI-TOF-MS. **e**, Potential silicate polymers in human tissue samples identified by MALDI-TOF-MS.

We analyzed insoluble silicon using MALDI-TOF-MS. Based on existing literature and silicon chemistry, insoluble silicon is expected to exist primarily as polysilicates (Fig. 2c). Synthetic silicate polymers (1% w/w), prepared by neutralizing sodium silicate, served as a standard for validation and calibration. Mouse tissues were enzymatically digested and washed to minimize interference from biomolecules and salts before MALDI-TOF-MS analysis. The results revealed that insoluble silicon in mouse tissues exhibited spectral characteristics identical to the synthetic silicate polymers standard (Fig. 2d and Fig. S2). Silicate polymers displayed a molecular weight distribution between 700-5000 Da, with characteristic peaks separated by 24 Da, corresponding to the monomeric silicic acid unit (Fig. 2c). This indicates a degree of polymerization (n) ranging from approximately 10-200. Mouse spleen, brain, bone marrow, and bone exhibited broader peaks and wider molecular weight distributions, suggesting greater diversity in polymerization, particularly in bone tissue, where the distribution shifted towards higher molecular weights.

Applying the same methodology, we identified insoluble silicon compounds in several human tissues (skeletal muscle, liver, stomach, colon, small intestine, and bone). Notably, human skeletal muscle and bone specimens contained silicate polymers analogous to those observed in murine tissues, characterized by molecular weights of 1301.275 Da and 1228.928 Da, yet exhibiting lower polymerization degrees compared to their murine counterparts (Fig. 2e). Surprisingly, characteristic molecular weight profiles of silicate polymers were undetectable in human hepatic, stomach, colonic, and small intestinal tissues, indicating the absence of typical insoluble silicon compounds dominated by silicate polymers in these human tissues. This finding further reveals substantial interspecies variations in the chemical speciation of silicon within biological tissues among mammalian species.

Soluble silicon species were analyzed using UPLC/Triple-TOF-MS. Lacking a standard silicon compound database, we compiled a silicate database based on literature, encompassing various silicon species, including silicate oligomers (silicon atoms < 10), sugar (fructose, ribose)-silicate complexes, and catecholamine (dopamine, epinephrine, norepinephrine)-silicate complexes (Fig. S3 and Fig. S4)^43–48^. This database includes molecular weights, chemical formulas, and structures. Following reported methods, silicate oligomers were synthesized by mixing sodium silicate (2 mM) and hydrochloric acid (pH adjusted to 7.4) and analyzed by UPLC/Triple-TOF-MS in both positive and negative ion modes. Matching against our database, we identified m/z values of 78.9842, 96.9946, 172.9567, 174.9730, 232.9229, 276.8953, and 484.8673, all within ±10 ppm of theoretical values, corresponding to polymerization degrees of 1 to 6 (excluding 5) (Table S2 and Table S3). And these oligomers eluted at approximately 0.02 minutes and 0.6-0.8 minutes. Applying this method to mouse tissues, we identified various silicate oligomers (n = 1, 3, 4, 5, 6, 7, 8) across all tissues (Table 2 and Table S3). Tetramers and hexamers, likely stabilized by six- or eight-membered ring structures, were most prevalent. These oligomers may exist in a metastable state, undergoing structural changes or polymerization under different physiological conditions. However, their inherent cyclic structures, or the presence of capping agents, may also confer stability and enable specific biological functions. Higher-order oligomers may exist as isomers with identical or distinct retention times, precluding definitive isomer identification. Notably, we identified organosilicon species in serum (fructose-silicate complex and tetracoordinate dopamine-silicate complex) and kidney (ribose-silicate complex). These organosilicon complexes, potentially exhibiting enhanced solubility and stability, may represent major forms of silicon transport and excretion.

**Table 2.**
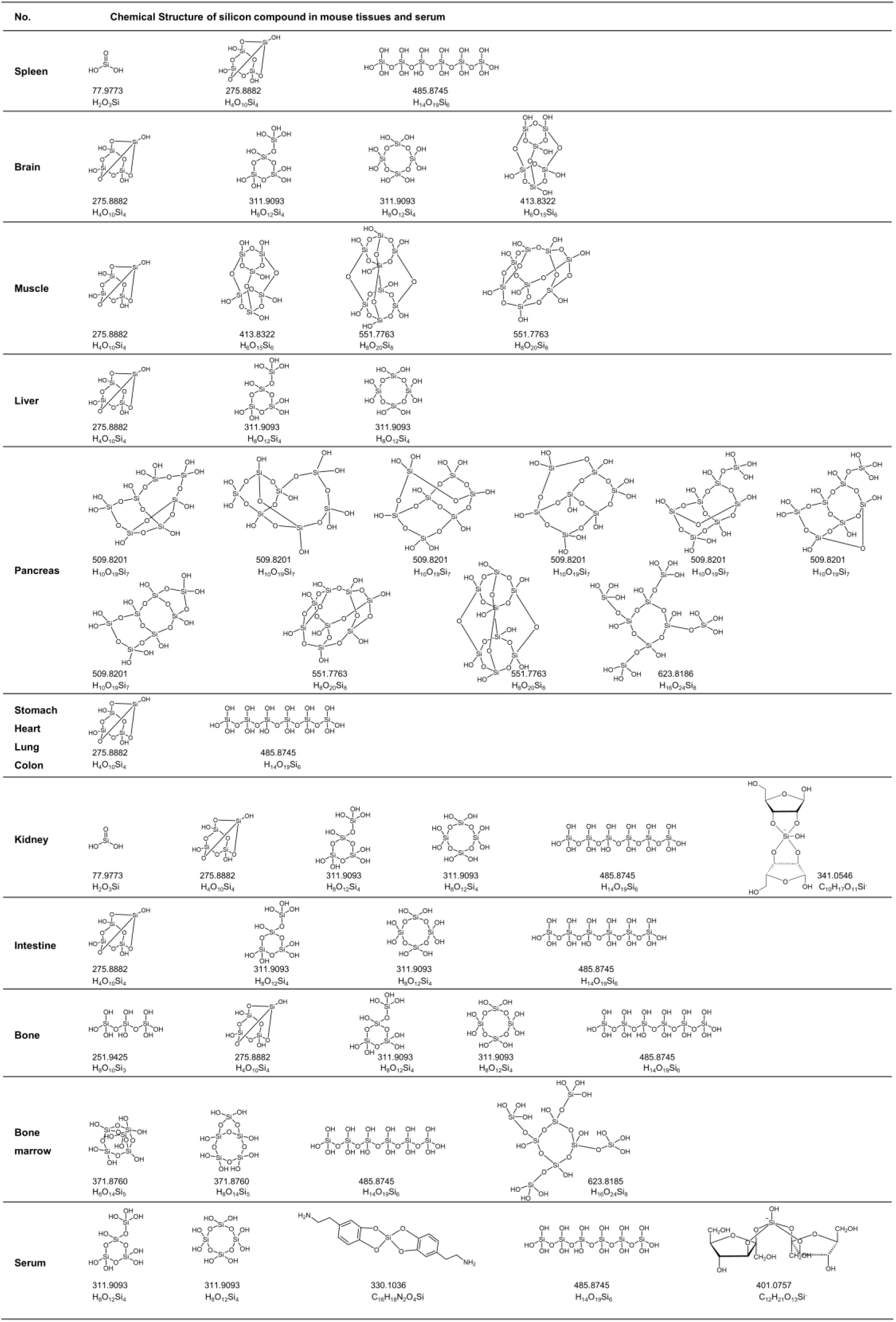
Chemical structure of potential silicon compounds in mouse tissues and serum identified by UPLC/Triple-TOF-MS.

Employing an identical analytical approach, we further investigated soluble silicon compounds in human tissues (skeletal muscle, liver, stomach, colon, small intestine, and bone). Similar to insoluble silicon compounds, the composition of soluble silicon compounds exhibited marked interspecies divergence between human and murine tissues. Notably, human tissues predominantly contained soluble silicate hexamers, heptamers, and nonamers, whereas the nonameric forms were undetectable in murine specimens. The linear hexameric silicic acid (MW = 485.8745) was universally detected across all human tissues, serving as the exclusive soluble silicon species in hepatic and osseous tissues. Intriguingly, this compound was absent in multiple murine tissues including brain, muscle, liver, and pancreas. Comparative analysis revealed greater diversity of organosilicon compounds in human tissues. Distinct organosilicon complexes were tissue-specifically identified: ribose-silicate complexes in muscular tissue, fructose-silicate complexes in colonic and intestinal tissues, norepinephrine-silicate complexes in gastric tissue, with dopamine-silicate and epinephrine-silicate complexes observed in circulation. These sophisticated organosilicon derivatives demonstrated enhanced solubility and stability compared to simple silicate polymers, suggesting their potential significance in human-specific biological functions (Table 3 and Table S4).

**Table 3.**
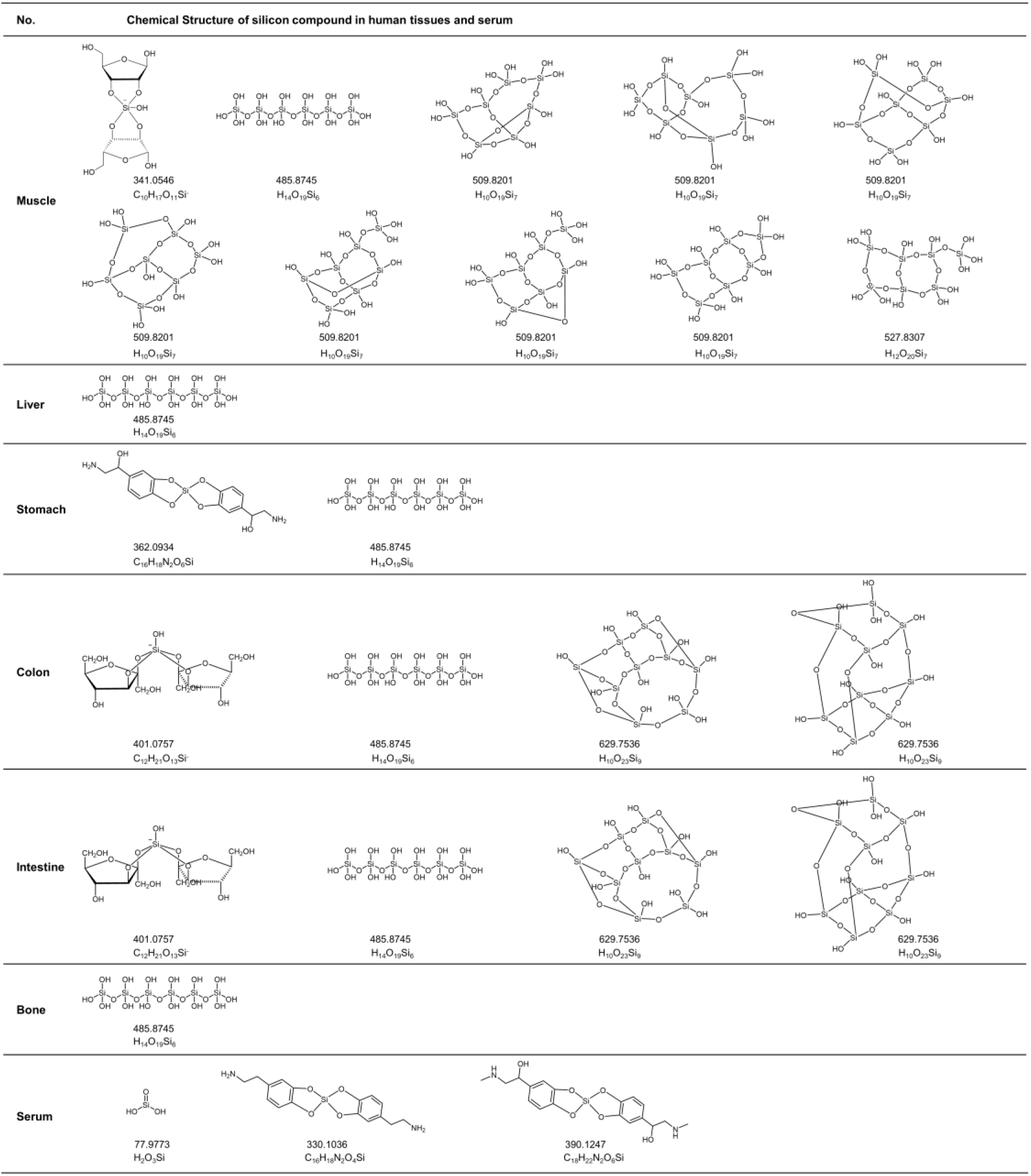
Chemical structure of potential silicon compounds in human tissues and serum identified by UPLC/Triple-TOF-MS.

In summary, our systematic characterization through two distinct mass spectrometric approaches has revealed the co-existence of insoluble silicon compounds (predominantly silicate polymers) and soluble silicon species (encompassing silicate oligomers and organosilicon derivatives) in both murine and human tissues. This comparative analysis demonstrated pronounced interspecies divergence in silicon speciation profiles between murine and human biological systems. The identification of diverse silicon-containing molecular architectures across examined tissues strongly suggests silicon’s potential involvement in specialized biological processes.

### Silicon Visualization in Animal Tissues *via* Micro Area X-ray Fluorescence Spectroscopy (micro-XRF)

The high abundance and molecular complexity of silicon compounds in murine and human tissues suggest that their intra-tissue distribution may exhibit unique patterns, potentially correlating with certain known tissue structures. Understanding the distribution characteristics of silicon compounds in animal tissues will provide critical insights into their biological functions. However, due to silicon compounds’ lack of specific biochemical chromogenic reactions and low immunogenicity, conventional histological staining techniques (histochemical or immunohistochemical staining) currently cannot visualize silicon’s spatial distribution.

Therefore, in this study, we attempted to employ micro-XRF for visualizing silicon distribution in murine and human tissues. Micro-XRF is an elemental imaging technique based on X-ray excitation-induced characteristic fluorescence spectra. By scanning the sample surface with a focused X-ray beam, it acquires elemental composition, concentration, and spatial distribution data, enabling elemental mapping through computational simulation. Given the paucity of prior micro-XRF studies on silicon in animal tissues, we established a silicon-specific imaging protocol by referencing existing literature on micro-XRF analysis of iron, copper, and chlorine in tissue sections (Fig. 3a). Briefly, adhering to micro-XRF technical requirements, we fixed tissues, embedded them in paraffin, sectioned the blocks using a microtome to expose smooth cutting surfaces, and performed micro-XRF scanning on these surfaces to obtain silicon distribution data.

**Figure 3.**
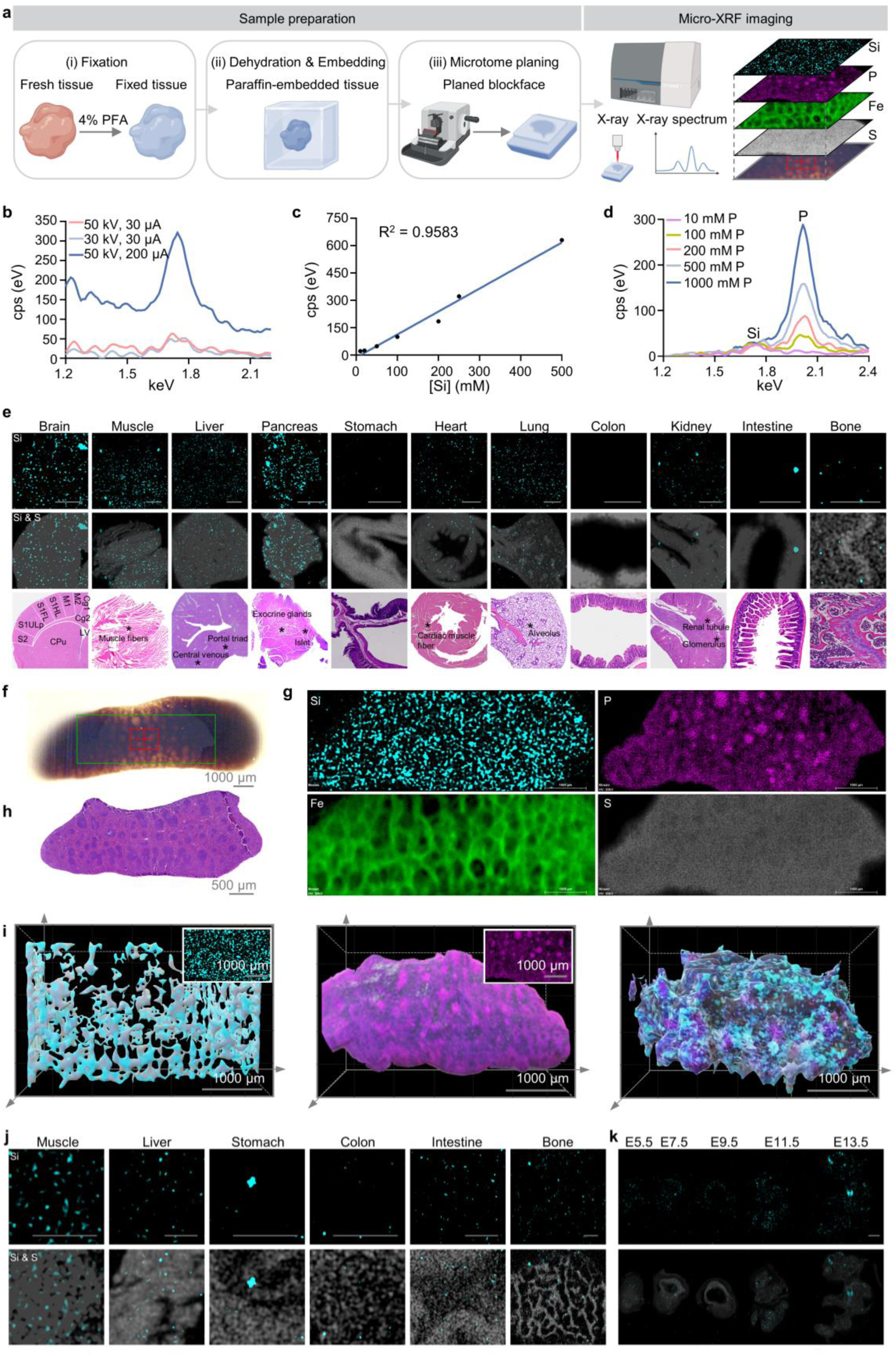
Silicon distributions in animal tissues are captured by the micro-XRF approach. **a**, Schematic of the micro-XRF experimental setup for silicon detection in tissue samples. The tissue samples are acquired using procedures including fixation, dehydration, paraffin embedding, and surface flatting. **b**, XRF spectra of a synthetic silicon-containing sample (50 mM sodium silicate) taken with various acquisition parameters (voltage and current). **c**, A calibration curve of the XRF silicon signals of various synthetic samples. There is a linear correlation between the concentrations of silicon in the synthetic samples and the XRF signals. **d**, Silicon peaks in XRF spectra of different concentrations of sodium phosphate encapsulated in synthetic samples containing 20 mM sodium silicate. **e**, Representative silicon mapping, merge mapping (silicon and sulfur), and histological staining of mouse tissues. Scale bars, 1000 μm (colon, 500 μm). **f-h**, Element mapping, and histological staining of mouse spleen. **i**, 3D reconstruction views of silicon, phosphorus, and merge mapping in the mouse spleen (Reconstructed by Tripo3d.ai; Scale bars, 1000 μm). **j**, Representative silicon mapping and merge mapping (silicon and sulfur) of human tissues. Scale bars, 1000 μm. **k**, Representative silicon mapping and merge mapping (silicon and sulfur) of mouse embryos. Scale bar, 1000 μm. All test samples are embedded in paraffin. All spectra are recorded by an X-ray beam with 50 kV high voltage and 30 μA current focused to 20 μm diameter spot. The sample is moved in a stepwise fashion in 5 μm increments with respect to the focused beam and X-ray fluorescence spectra are collected for 4 ms at each pixel.

First, we needed to confirm the X-ray spectral characteristics of silicon under paraffin-embedding conditions, determine the spectroscopic properties of silicon in our analytical workflow, select appropriate parameters, and evaluate potential interferences. To this end, we prepared sodium silicate-dextran lyophilized composite materials as reference samples, paraffin-embedded them (Fig. S5a), and subjected them to micro-XRF analysis. We utilized these samples with known silicon content to calibrate the system, focusing on analytical specificity and sensitivity within defined concentration ranges. Results demonstrated that exposed surfaces of synthetic samples effectively responded to micro-XRF silicon scanning and data acquisition. Using these simulated tissue samples, we identified silicon’s characteristic X-ray fluorescence peaks within the 1.6–1.9 eV range. Additionally, we observed that XRF sensitivity increased with higher photon flux at elevated energies and reduced currents (Fig. 3b). Consequently, we adopted 50 kV voltage and 30 μA current for all samples to optimize sensitivity and minimize noise. Subsequent testing of synthetic materials with varying silicon concentrations (10–500 mM) revealed a linear correlation between silicon concentration and XRF signal intensity (Fig. 3c and Fig. S5b), confirming the protocol’s accuracy. We specifically evaluated phosphorus interference by adding sodium phosphate (10–1000 mM) to synthetic samples, finding no significant cross-impact on silicon analysis (Fig. 3d and Fig. S5c).

Next, to apply micro-XRF for mapping elemental distributions in specific tissue regions, we further optimized imaging parameters including scan step size, time per pixel, and contrast. Preliminary experiments using synthetic materials determined optimal settings: 5 μm scan step size and 4 ms dwell time per pixel. These parameters balanced signal intensity, noise suppression, and imaging duration, enhancing resolution and signal-to-noise ratio. During image processing, we adjusted contrast and removed paraffin background noise to ensure clarity and accuracy of silicon distribution maps. Applying these parameters to biological tissues, micro-XRF-generated silicon heatmaps exhibited high resolution and fidelity, with randomly selected signal points matching the spectral characteristics of synthetic reference samples. Further analysis revealed significant heterogeneity in silicon distribution across tissues (e.g., brain, heart, muscle, liver, spleen, bone), ranging from near-noise baseline levels to highly enriched regions (Fig. S6). Thus, paraffin embedding combined with optimized micro-XRF enabled high-resolution, precise visualization of macroscopic silicon distribution patterns in tissues.

Following methodological optimization, we systematically analyzed silicon distribution in multiple murine tissues via micro-XRF (Fig. 3e). Generally, silicon exhibited clustered distribution patterns across murine tissues, forming localized hotspots. These hotspots appeared uniformly dispersed within tissues, showing no correlation with established histological structures (e.g., parenchymal cell distributions, extracellular matrix organization, or tissue-specific laminar architectures). Silicon compounds instead appeared to exist as distinct independent structures within murine tissues. Given sulfur’s high and uniform distribution across proteins, concurrent sulfur mapping *via* micro-XRF outlined tissue architecture. Co-localization analysis of sulfur and silicon in identical regions facilitated deeper investigation of silicon distribution patterns. For enhanced accuracy, we cross-referenced results with H&E-stained sections of corresponding tissues.

Our analysis revealed pronounced silicon clustering in low-structural-intensity tissues like brain and pancreas, with larger cluster coverage. In murine prefrontal cortex (Cg1, Cg2), silicon distribution was more abundant, whereas in somatosensory/motor cortices and gray matter regions (S1, S1ULp, S1FL, M1, M2, CPu), silicon followed specific topological patterns aligned with neural network nodes. Pancreatic silicon localized to exocrine interstitial spaces rather than islets. In liver, kidney, and lung tissues, silicon dispersed uniformly within extracellular matrices without spatial correlation to cells or structural landmarks. For instance, pulmonary silicon distribution did not align with alveolar or acinar structures. In muscle and heart tissues with ordered myofiber arrangements, silicon adopted speckled distributions within intermuscular gaps rather than tracking fiber orientation. Murine femoral silicon concentrated in cancellous bone lacunae rather than trabeculae (gray-white regions). Rat femoral coronal sections more clearly demonstrated silicon enrichment in marrow versus trabeculae (Fig. S7a), contradicting prior assumptions about silicon’s bone-strengthening role and suggesting potential involvement in hematopoietic functions.

Murine spleen yielded the clearest micro-XRF images, likely due to minimal extracellular matrix interference. High-resolution spleen mapping (Figs. 3f-3h) revealed interconnected silicon clusters forming a three-dimensional network—a feature potentially present but undetectable in other tissues. As current micro-XRF systems lack Z-axis reconstruction capability, we employed Tripo AI for 3D simulation of Fig. 3f regions, approximating this silicon framework (Fig. 3i). Concurrent phosphorus mapping in spleen (Fig. 3g) demonstrated silicon domains enveloping phosphorus-marked cellular regions, consistent with extracellular matrix localization observed in Figs. 1j-1l. We hypothesize similar silicon networks exist in other tissues but remain obscured by dense extracellular matrices, manifesting as discrete clusters in sectional scans. Human skeletal muscle, liver, stomach, colon, small intestine, and bone analyses revealed analogous silicon distribution patterns to murine tissues but with larger clusters (e.g., skeletal muscle). Strikingly, human bone exhibited trabecular silicon enrichment (Fig. 3j), contrasting sharply with murine/rat observations.

Embryonic silicon distribution analysis (Fig. 3k) showed uniform silicon distribution across epiblast (EPI) and extraembryonic ectoderm (ExE) during early gastrulation (E5.5). As development progressed (E7.5), silicon gradients emerged with germ layer differentiation—mesoderm-derived tissues (heart, muscle, bone) showed progressive enrichment, while endoderm-derived organs (lung, liver, pancreas, intestine) and ectoderm-derived structures (nervous system, epidermis) exhibited tissue-specific localization. Organogenesis stages (E9.5, E11.5, E13.5) demonstrated regionally restricted silicon accumulation, particularly in developing bone. These spatiotemporal patterns suggest silicon’s involvement in critical developmental processes. Silicon detection in avian yolk sacs (Fig. S7b) further implies evolutionarily conserved biological roles. Given reported teratogenic effects of silicon deficiency on limb patterning^17,18^, we hypothesize silicon compounds participate in morphogenesis and organogenesis.

Together, silicon compounds exhibit unique distribution patterns in animal tissues, forming an architecturally independent spatial network distinct from protein-based structures. This network may directly interact with cellular components and participate in diverse biological processes.

## Discussion

Although some studies have indicated that silicon deficiency leads to developmental abnormalities in animals, silicon continues to be classified as a non-essential element for humans in many scientific literatures. This conclusion primarily stems from two longstanding perceptions: first, previous research reports have asserted that silicon content in human and animal tissues is extremely low (with approximately 1g of total silicon in a normal human body), and second, no studies have definitively identified the chemical structures of silicon compounds *in vivo* or clarified their participation in specific biological processes. Through methodological innovations, our study has accurately measured silicon content in mouse and human tissues, revealing that the actual silicon levels in animal tissues are 1-2 orders of magnitude higher than traditionally recognized. Based on this discovery, the total silicon content in a normal human body can be estimated to exceed 10 grams. This quantity suggests that silicon abundance in humans may approach that of magnesium, indicating that silicon should no longer be categorized as a “trace element” but rather as a major constituent in the human body, thereby implying its critical biological roles. Furthermore, we identified complex, tissue- and species-specific chemical structures of silicon in mouse and human tissues, as well as unique distribution patterns. These findings not only reinforce the likelihood of silicon compounds possessing definitive and essential biological functions but also establish foundational information and methodologically valuable approaches for future investigations into silicon’s biological activities.

ICP-OES is the most widely used elemental analysis technique, offering simplicity and high precision for most elements, particularly metals. However, its applicability is restricted for certain elements due to limitations in sample pretreatment protocols, which prohibit the use of strong alkalis or hydrofluoric acid. Historically, nearly all studies have employed ICP-OES for silicon quantification in animal tissues, driven by the entrenched assumption that silicon exists at negligible concentrations *in vivo*. This presumption led researchers to overlook the potential impact of strong acids on low-concentration silicon compounds. Paradoxically, this very assumption originated from early ICP-OES-based silicon quantifications. Our critical analysis of the conventional acid digestion-ICP-OES method revealed that the acid digestion process induces severe polymerization and precipitation of silicon compounds, rendering ICP-OES fundamentally unsuitable for accurate quantification. In contrast, the molybdenum blue colorimetric method—a well-established silicon quantification technique predating ICP-OES—permits flexible sample pretreatment strategies to ensure complete silicon extraction. This method is universally adopted as the standard for silicon analysis by regulatory agencies in many countries and is routinely applied to quantify silicon in plant tissues^7,12,15,27–32^. After optimizing pretreatment protocols, we successfully adapted this method for animal and human tissues, achieving exceptional precision.

Elucidating the chemical nature of silicon is pivotal for uncovering its biological functions. However, research on silicon speciation in animal systems remains scarce due to technological limitations. By integrating silicon’s intrinsic properties, silicon compound molecular libraries, and mass spectrometry data, we identified a series of silicate oligomers and polymers in mouse and human tissues. This breakthrough provides the first clear characterization of biologically derived silicon compounds in animal tissues. Nevertheless, several limitations require future refinement: First, the current database of soluble silicon compounds remains incomplete. Expanding the reference library with additional naturally occurring silicon compounds and enriching it with molecular formulae, molecular weights, structural details, chemical properties, and spectral data (e.g., mass spectra, chromatographic profiles, nuclear magnetic resonance, and infrared spectra) will enhance analytical robustness. Second, the inability to quantify specific silicon compound molecules hinders precise identification of their roles in biological functions. Introducing certified reference materials or stable isotope labeling strategies during mass spectrometry could enable accurate measurement of silicon compound levels in specific biological contexts^49^. Third, silicon extraction protocols may alter the polymerization degree and molecular structure of silicon compounds. Optimizing enrichment and extraction processes to preserve their native chemical characteristics remains imperative. Fourth, the extensive presence of silicate oligomer isomers poses challenges for their differentiation, purification, and subsequent spectral analysis (e.g., NMR). Addressing these challenges will require innovative separation and enrichment methodologies.

Silicon imaging in animal tissues offers critical insights into its spatial distribution. However, the chemical inertness and low immunogenicity of silicon compounds have impeded the development of conventional imaging probes (e.g., fluorescent tags or antibodies). Our study employed micro-XRF elemental imaging, which utilizes X-ray fluorescence emitted upon excitation of outer-shell electrons to map elemental distributions. This approach confirmed silicon’s widespread and tissue-specific presence in animal tissues. Nevertheless, its limited spatial resolution restricts detailed analysis. Future advancements should focus on: 1) Enhancing resolution to cellular or subcellular levels using techniques like synchrotron radiation X-ray fluorescence (SXRF) or nano-scale secondary ion mass spectrometry (Nano-SIMS)^50–52^; 2) Coupling elemental imaging with exogenous element-labeled probes or antibodies to co-localize silicon with biomolecules (nucleic acids, proteins, lipids, or carbohydrates), thereby elucidating silicon-biomolecule interactions; and 3) Developing multimodal imaging strategies. For specific silicon compounds, two parallel approaches could be pursued: designing high-affinity molecular probes (e.g., fluorescent probes or aptamers) and integrating mass spectrometry imaging (e.g., DESI-MSI, MALDI-MSI) with spatial omics for spatiotemporal analysis of silicon compound dynamics and interactions^53^. Synergizing these technologies will provide comprehensive, complementary insights into silicon’s spatiotemporal behavior *in vivo*.

By refining and developing analytical methods for silicon in animal tissues, we made the following key discoveries: Similar to plants, silicon exists at elevated concentrations in animals, adopts unique molecular architectures, and exhibits tissue-specific distribution patterns. Moreover, silicon content, speciation, and distribution vary significantly across species. These findings position silicon as an essential element with critical physiological roles in animals and establish a foundation for future functional studies. However, comprehensive understanding of silicon’s biological functions demands resolution of several outstanding questions: 1) Clarifying silicon absorption and transport mechanisms; 2) Identifying target cells, receptors, and molecular pathways; 3) Validating biological functions through cellular and animal models; and 4) Investigating silicon’s roles in physiological and pathological processes across species. Addressing these questions will require continued methodological advancements. For instance, characterizing silicon speciation may reveal whether highly soluble organic silicon compounds mediate its systemic circulation. Identifying tissue-specific silicon compounds and combining them with co-immunoprecipitation could uncover molecular targets and mechanisms. Systematic optimization of silicon detection methodologies remains indispensable for resolving these fundamental biological questions.

## Methods

### Reagents and resource

All solutions were prepared using ultra-high purity (UHP) water (18 MΩ/cm) from a Millipore water purifier. Polypropylene plastic tubes (JET Biofil), rinsed with UHP water, were used throughout. All chemicals were of high purity (Sigma-Aldrich or Aladdin). All food samples were purchased from the market (Qixia, Nanjing, China), transported to the laboratory, and thoroughly rinsed with UHP water to remove soil and other surface contaminants. Study protocols involved human samples collection and treatment were approved by Affiliated Hospital of Jiangnan University (ethical approval number: LS2022109). Written informed consent was obtained from each individual before enrolment. All procedures complied with relevant ethical regulations. The surgically resected normal adjacent tissues were quickly frozen in liquid nitrogen in the polypropylene plastic tubes until analysis or immediately prepared for examination.

### Animals

All mouse care and procedures were performed in accordance with animal experimental protocols approved by Nanjing University Institutional Animal Care and Use Committee (ethical approval number: IACUC-2006007). The wild-type C57BL/6J mice were purchased from Vital River Laboratories (Beijing, China), fed sufficient food and water, and maintained on a regular 12-hour day/night cycle under specific pathogen-free conditions.

### Tissue digestion

All samples were dried by a lyophilizer for 2 days prior to analysis. The dried sample (tissues, around 20 mg; serum, 100 μL) was then acid-digested in a mixture of 0.5 mL of HNO_3_, 0.5 mL of H_2_O_2_, and 0.5 mL of HCl at 95°C for 3 h until the digestion solution became transparent and clear. Then concentrate the solution to about 1 mL, and transfer 150 μL quantitatively to a polypropylene tube. The Si level in the digest solution of the tube was determined as described below. Just prior to the colorimetric determination, adjust the acidic solution to approximate neutrality, using 1.0 N sodium hydroxide and 0.1 N hydrochloric acid.

### Silicon determination

The silicon level in the digest solution was determined by the modified colorimetric molybdenum blue method. In a polypropylene tube, the pH of 150 μL of the digest solution (except for the 5 μL of bone digest solution and 50 μL of liver or muscle digest solution), containing around 3 mg dry tissue, was initially adjusted to approximately 7.0, followed by the addition of ultrapure water to achieve a final volume of 2 mL. And then the solution was treated with 0.4 mL of HCl (150 g/L), 0.1 mL of NaF (20 g/L) and agitated for a while to fully dissolve silicon. In the presence of HCl, any form of polysilicon is depolymerized by treatment with HF. Then the mixture solution was added under stirring by 2 mL of boric acid (48 g/L) as acid binding agent. This was followed by the addition of 1 mL of (NH_4_)_6_Mo_7_O_2_ (140 g/L) and the mixture were stirred for 5 minutes. The interference of phosphorus was removed by 0.5 mL of oxalic acid (100 g/L) and 3 mL of sulfuric acid (400 g/L). The resultant silico-molybdate complex was reduced by 0.2 mL of ascorbic acid (25 g/L), and the test solution was blue and transparent. Then add water to unify the volume of the solution to 10 mL. After 20 min, the absorbance was measured at 800 nm with a UV-spectrophotometer (TU-1800). The silicon level in the digest solution was also determined by inductively coupled plasma optical emission spectrometry (ICP-OES; PerkinElmer; Avio 500). Analytical line used was 251.2 nm.

### Spike and recovery experiment

The albumin concentration in serum is approximately 0.6 mM, while in the serum test solution, it ranges around 0.06 mM. In mouse bone tissue, phosphorus content is approximately 15%-20%, with phosphorus concentrations in the bone test solution ranging from 3 to 4 mM. To demonstrate that the HF-MB method can accurately detect silicon in animal tissues without interference from proteins or phosphorus, we performed a dilution linearity test using albumin/disodium hydrogen phosphate mixtures (BSA 0.55 mM, Na_2_HPO_4_ 4.31 mM, 150 μL digestion solution), or mouse tissue samples (spleen, heart, and lung) (3 mg dry tissue, 150 μL digest solution) spiked with Si standard solution. Samples were assessed by adding 150 μL of synthetic or mouse tissue sample (acid digestion solution) and 10 μL of spike Si standard solution (2-40 μg) calculated to yield the intended spike levels. The reported values for spiked samples were calculated by subtracting the endogenous (no-spike) value. The recovery rates for the spiked samples were determined by subtracting the value of the no-spike sample from the spiked sample, then dividing the resulting difference by the theoretical value of the added standard.

### Mouse tissues decellularization

To obtain the whole-tissue decellularized ECM from mice, the tissues were decellularized following an in-situ perfusion protocol. Briefly, mouse was anaesthetized with 4% chloral hydrate and intraperitoneally injected with heparin sodium (35 mg/mL, 300 μL). The left ventricle was cannulated with a polyethylene tube attached to a peristaltic pump, and the right atrium was severed to perfuse out the buffer. For liver, the hepatic portal vein was cannulated with a polyethylene tube attached to a peristaltic pump, the inferior vena cava was severed to perfuse out the buffer. Heparin sodium solution (1 mg/mL) was perfused for 1 h to flush out blood. Then 0.1% trypsin, 0.05% EDTA and 1% SDS solutions were sequentially perfused for 3 h. Then 1% triton solution was perfused to until the tissue was translucent and the deionized water was perfused for 2 h to thoroughly rinse the buffer residue in decellularized ECM.

### Cell samples isolated from mouse tissues

To prepare cell samples, mouse heart was dissected into fragments using clean experimental scissors, digested with a protease cocktail (collagenase II and IV, 10 mg/mL each, WARBIO) at 37°C for 30 minutes, and then gently dissociated using 70-μm cell strainers. For liver tissue, mouse liver was isolated and digested with collagenase IV (1.64 mg/mL) and deoxyribonuclease I (1 mg/mL, Yeasen), followed by dissociation through 70-μm cell strainers. For spleen tissue, mouse spleen was fragmented and digested with 5 mL of accutase (STEMCELL Technologies) at 37°C for 60 minutes, followed by dissociation through 70-μm cell strainers. For lung tissue, mouse lung was dissected and digested with a protease cocktail (collagenase II, 1 mg/mL; collagenase I, 0.5 mg/mL, WARBIO; deoxyribonuclease I, 0.1 mg/mL) at 37°C for 40 minutes, and then gently dissociated using 70-μm cell strainers. For kidney, mouse kidney was dissected, digested with 8 mL of collagenase II (1 mg/mL) at 37°C for 20 minutes, and gently dissociated under 70-μm cell strainers. After collection, the cell suspensions were centrifuged at 300 g for 5 minutes at 4°C. The supernatant was removed, and the resulting cell pellets were obtained for further analysis.

### Quantitation assay of the hydroxyproline and glycosaminoglycan in tissues

The contents of hydroxyproline in tissues were measured using a hydroxyproline assay kit (Nanjing Jiancheng Technology Co.; A030-2-1) to quantify collagen content following the manufacture’s protocol. The data are expressed as hydroxyproline (in micrograms) / tissue wet weight (in grams). The contents of glycosaminoglycan in tissues were measured using a glycosaminoglycan kit (Nanjing Hanzhang Biotechnology Co.; HL19239.2).

### Preparation of mouse and human plasma and tissues for UPLC/Triple-TOF-MS analysis

Plasma (50 μL) was mixed with 150 μL of a 2:1 mixture of acetonitrile and methanol, followed by vortexing for 30 s. The mixture was then centrifuged at 13,000 rpm for 10 min at 4°C, and the supernatant was transferred to a UPLC/Triple-TOF-MS (AB Sciex, Triple TOF 4600) vial.

For tissue samples, 200 mg of tissues were homogenized in 500 μL PBS solution using a homogenizer. The resulting tissue homogenates were centrifuged at 13,000 rpm for 10 min at 4°C. The soluble fractions (100 μL) were mixed with 300 μL of a 2:1 mixture of acetonitrile and methanol. The mixture was then centrifuged at 13,000 rpm for 10 min at 4°C, and the supernatant was subsequently transferred to a UPLC/Triple-TOF-MS vial.

### Preparation of mouse and human tissues for MALDI-TOF-MS analysis

In this procedure, 25 μL of proteinase K (Takara) in 1 mL of TNES buffer (50 mM Tris, pH 7.5; 400 mM NaCl; 20 mM EDTA; 0.5 % SDS) was added to each insoluble tissue sample (100 mg). The mixture was vortexed briefly and incubated in a water bath at 57 °C for 8 h. The proteolysis was terminated by adding 1*PBS to wash out the proteinase solution. The sample was then centrifuged at 13,000 rpm for 10 min at 4°C to collect the insoluble components. The washing process was repeated twice. Subsequently, the insoluble components were treated with glycoside hydrolases (0.1% Amyl glucosidase, 1% β-galactosidase at 57 °C for 24-36 h), deoxyribonuclease I (1% at 37 °C for 8 h), and lipase (0.1% at 37 °C for 8 h) to remove biomolecules such as polysaccharides, nucleic acids, and lipids. The final mixture was centrifuged again at 13,000 rpm for 10 min at 4°C to obtain the insoluble components. To determine the mass of silicate polymers in the insoluble components, matrix-assisted laser desorption ionization time-of-flight mass spectrometry (MALDI-TOF-MS, Bruker Daltonics, UltrafleXtreme) analysis was performed. Dithranol (0.1 M) in tetrahydrofuran was used as the matrix, with silver trifluoroacetate (5 mg/mL) serving as the cationizing agent.

### Synthetic oligomeric and polymeric silicate samples

Different concentrations of sodium silicate solutions were prepared for oligomers (2 mM sodium silicate) and polymers (1%(w/w) sodium silicate), with the pH adjusted to 7.4 using HCl.

### Sample preparation for micro-XRF analysis

Tissue samples were first fixed with 4% paraformaldehyde and then dehydrated with different gradients of ethanol. After that, samples were fixed in paraffin blocks and cut the surface flat. Micro-XRF analyses were conducted at the high-performance micro area X ray fluorescence spectrometer (Bruker/M4 Tornado) using X-ray beam with 50 kV high voltage focused to 20 μm diameter spot. The sample was moved in a stepwise fashion in 5 μm increments with respect to the focused beam and X-ray fluorescence spectra were collected for 4 ms at each pixel.

### H&E staining

Tissues were fixed in 4% paraformaldehyde (PFA), processed, and embedded in paraffin. The sections (5 μm thick) were prepared and stained according to the manufacturer’s instructions (Jiancheng Bioengineering Institute, Nanjing, China).

### 3D reconstruction

The feature extraction and stereo-matching of silicon mapping within the tissues by micro-XRF were performed and transformed simultaneously with AI-powered 3D-modeling (www.tripo3d.ai).

### Statistical analysis

Statistical analysis was performed with SPSS (version 25.0) or GraphPad Prism (version 8.0) software. All values in graphs are presented as mean±SEM.

## Acknowledgments

This work was funded by the National Natural Science Foundation of China (32230056, 32361163656), Natural Science Foundation of Jiangsu Province (BK20232018), and University of Macau Research Committee (MYRG2023-00136-ICMS-UMDF).

## Author contributions

L.D. designed the study. Y.J.W. performed experiments and data analysis. L.D., Y.J.W., and Z.Y.L. provided guidance for the experiments and wrote the manuscript. R.Y.Y., G.X.Z., Y.L.W., C.Y.L., W.L.M., S.Y.L., Z.F.W., Q.Y.W., Z.X., and C.M.W. provided guidance for some experiments. Y.Y., and R.W. provided support for human samples.

## Data and materials availability

All data are available in the manuscript or the supplementary information.

## Supplementary information

**Figure S1.**
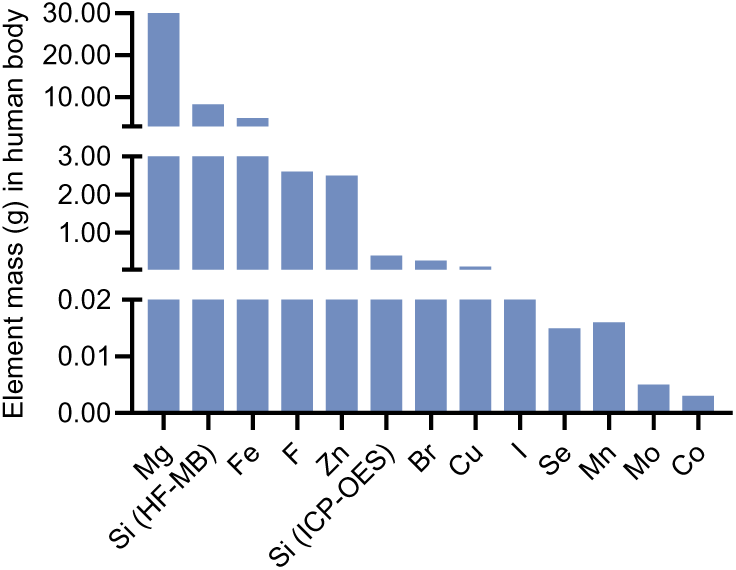
Related to Figure 1. Element mass in the human body (70 kg). Elements that constitute more than 0.01% of the human body are known as major elements, including C, H, O, N, K, S, Na, Cl, Mg, Ca, and P. Those that make up less than 0.01% are referred to as trace elements, including Si, Fe, F, Zn, Br, Cu, I, Se, Mn, Mo, Co. The trace element mass values are sourced from literature reports, whereas the silicon mass value is based on measurements conducted in this study.

**Figure S2.**
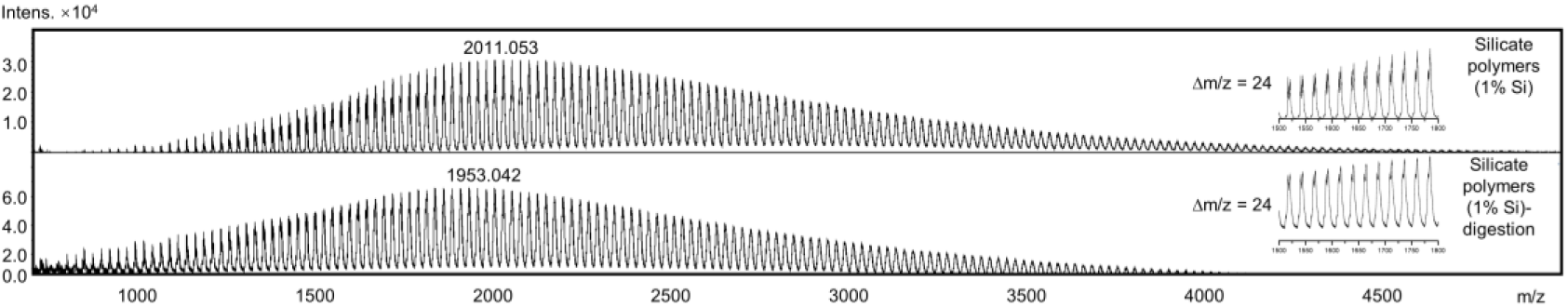
Related to Figure 2. Potential silicate polymers in synthetic silicate polymers or after enzyme digestion identified by MALDI-TOF-MS.

**Figure S3.**
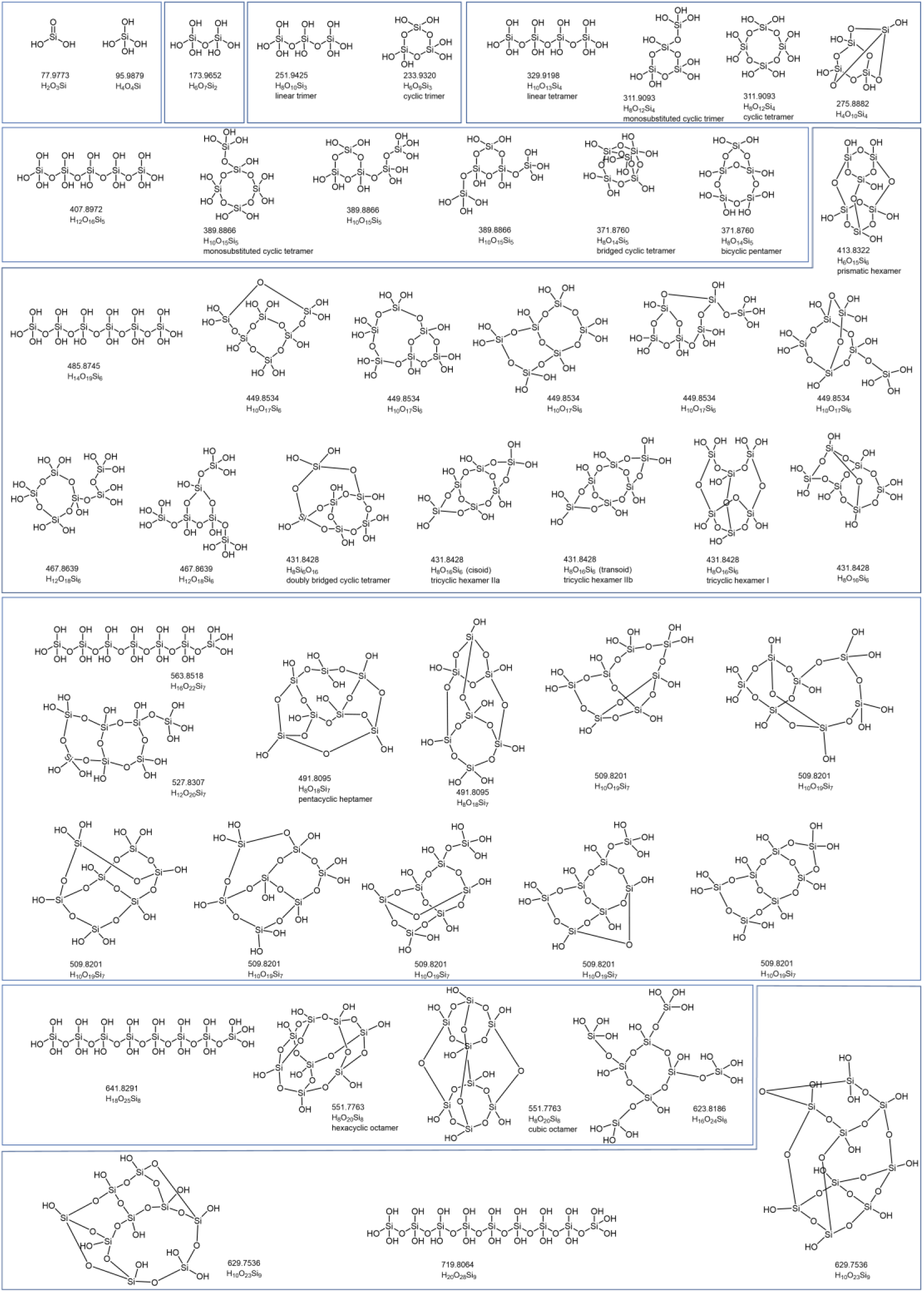
Related to Figure 2. Chemical structural formula library of silicate oligomers.

**Figure S4.**
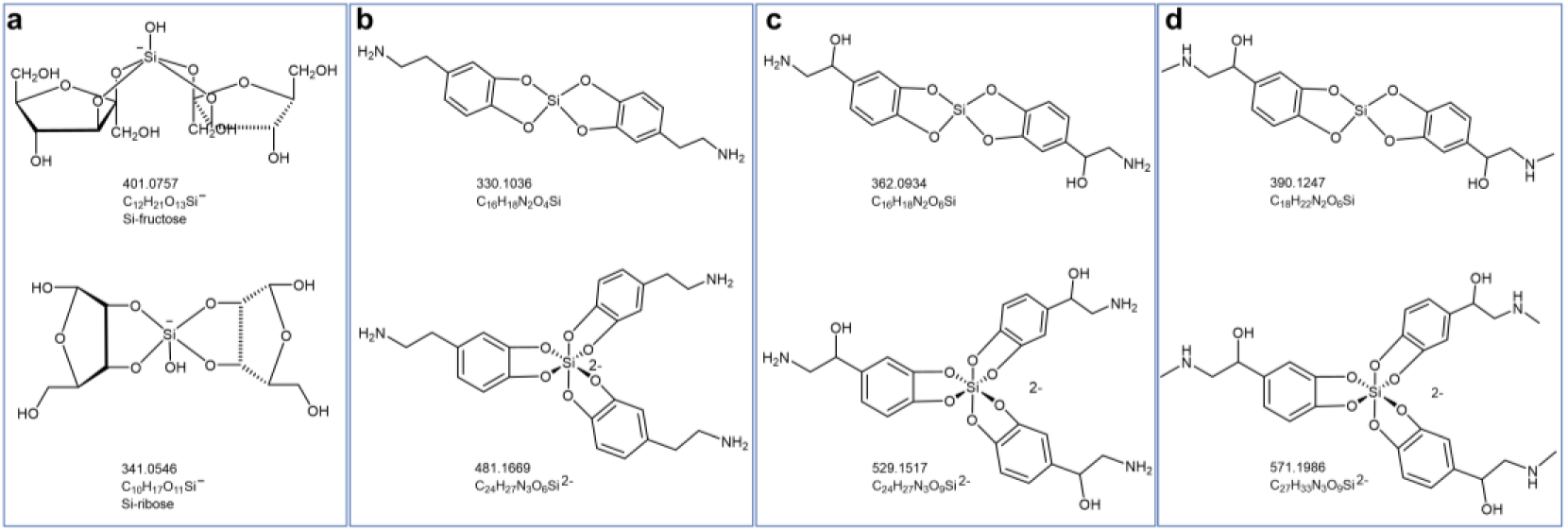
Related to Figure 2. Chemical structural formula library of sugar silicates (**a**), dopamine-silicates (**b**), norepinephrine-silicates (**c**), and epinephrine-silicates (**d**).

**Figure S5.**
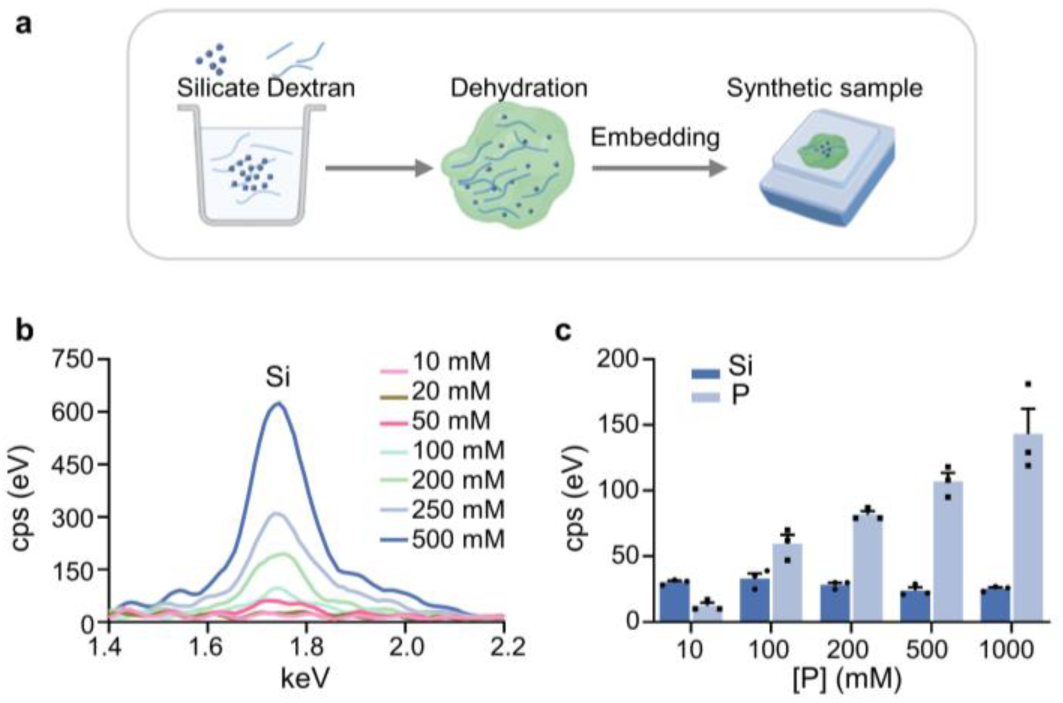
Related to Figure 3. **a**, Schematic of the micro-XRF experimental setup for silicon detection in synthetic samples. 100 mg of dextran powder is dissolved in 1 mL of sodium silicate solution, then lyophilized, embedded in paraffin and cut the surface flat as the synthetic samples. **b**, Silicon peaks in XRF spectra of different concentrations of sodium silicate encapsulated in synthetic samples. **c**, Silicon peak intensity in XRF spectra of different concentrations of sodium phosphate encapsulated in synthetic samples containing 20 mM sodium silicate.

**Figure S6.**
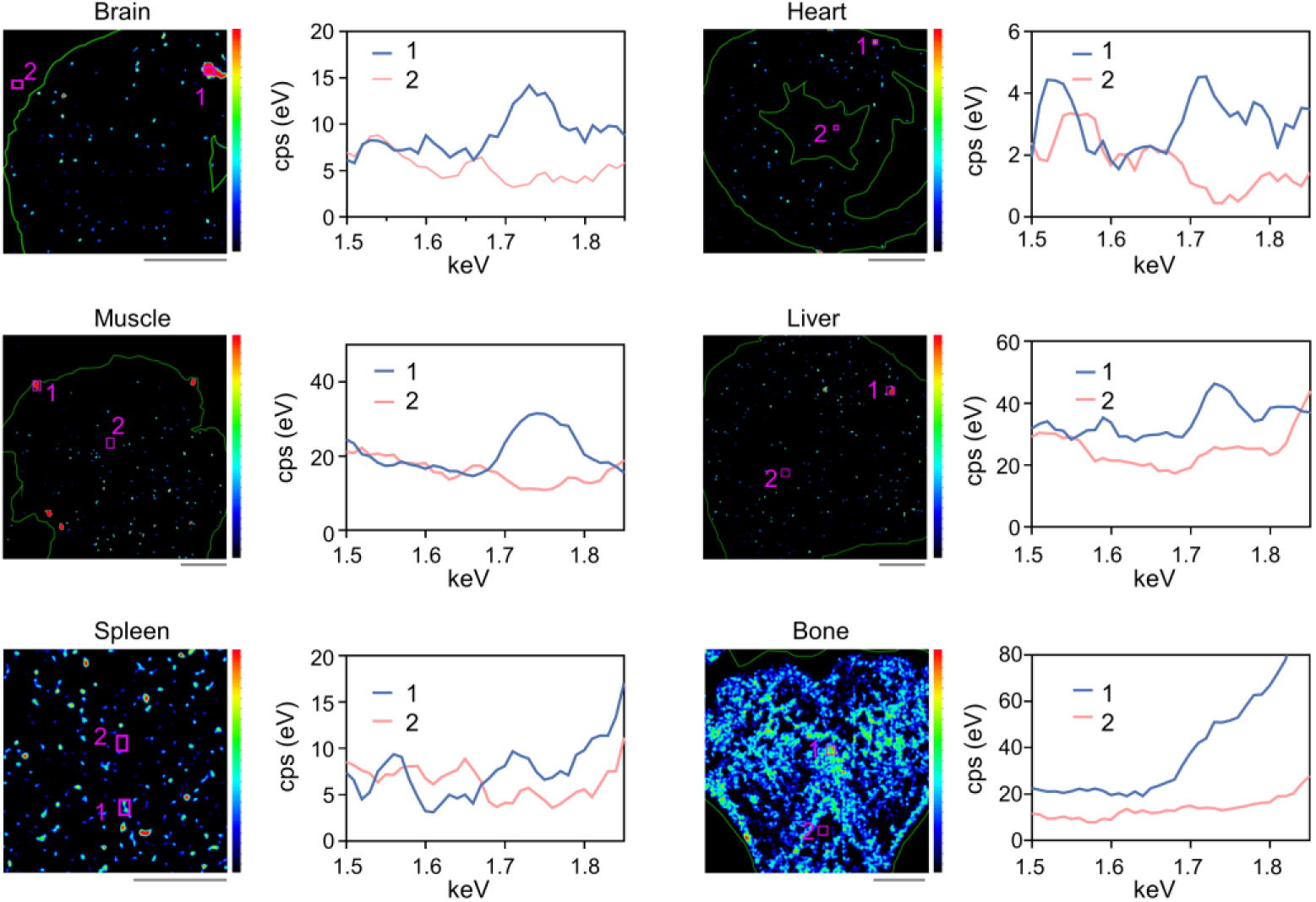
Related to Figure 3. Micro-XRF spectra (right) showing the silicon peaks, taken from the areas indicated by pink boxes in the micro-XRF heat maps (left) of different mouse tissues (brain, heart, muscle, liver, spleen, and rat bone). Scale bars, 1000 μm.

**Figure S7.**
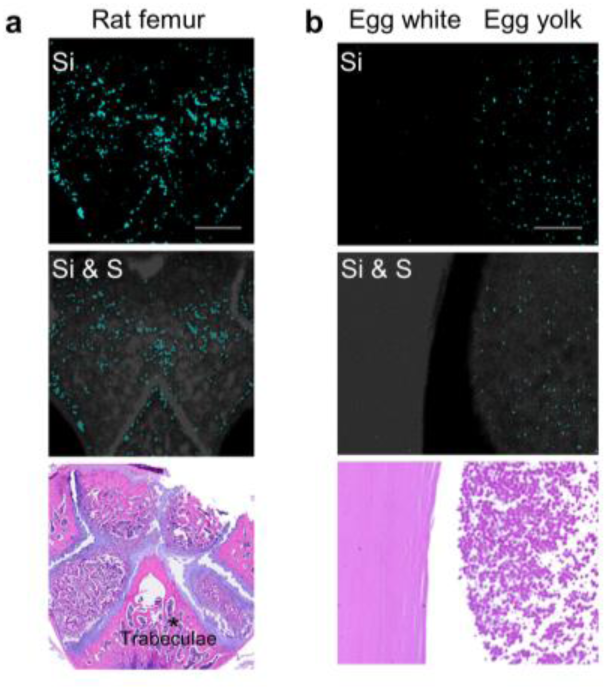
Related to Figure 3. Micro-XRF images are collected from rat femur, egg white and yolk. Scale bars, 1000 μm.

**Table S1.**
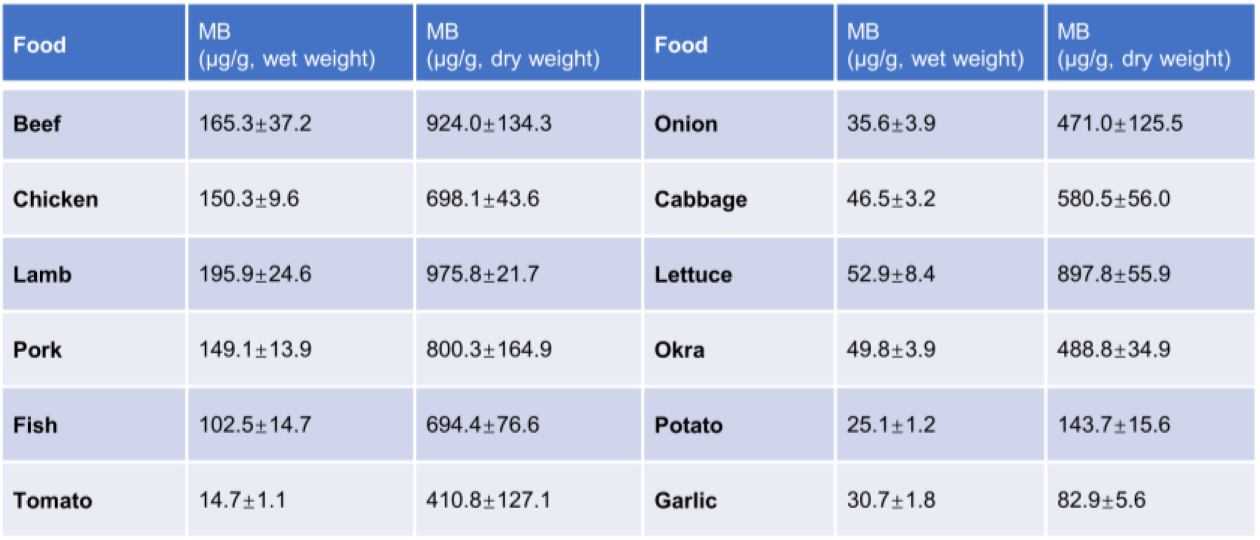
Related to Figure 1. Quantification of silicon in food determined by HF-MB method (n = 3).

**Table S2.**
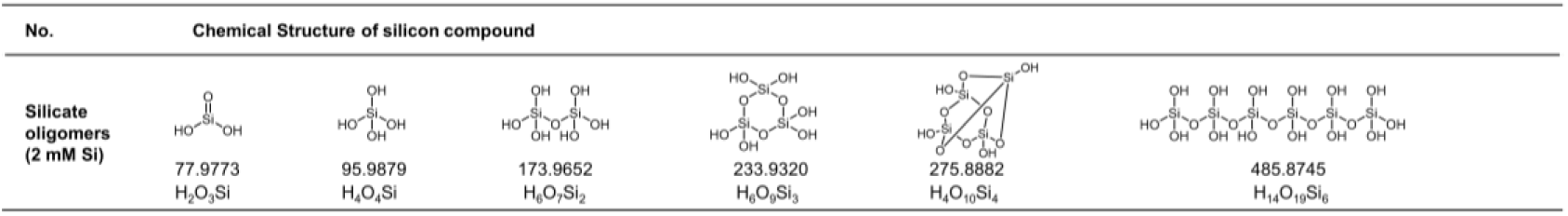
Related to Table 2. Chemical structure of potential silicon compounds in synthetic silicate oligomers solution (2 mM sodium silicate solution) identified by UPLC/Triple-TOF-MS.

**Table S3.**
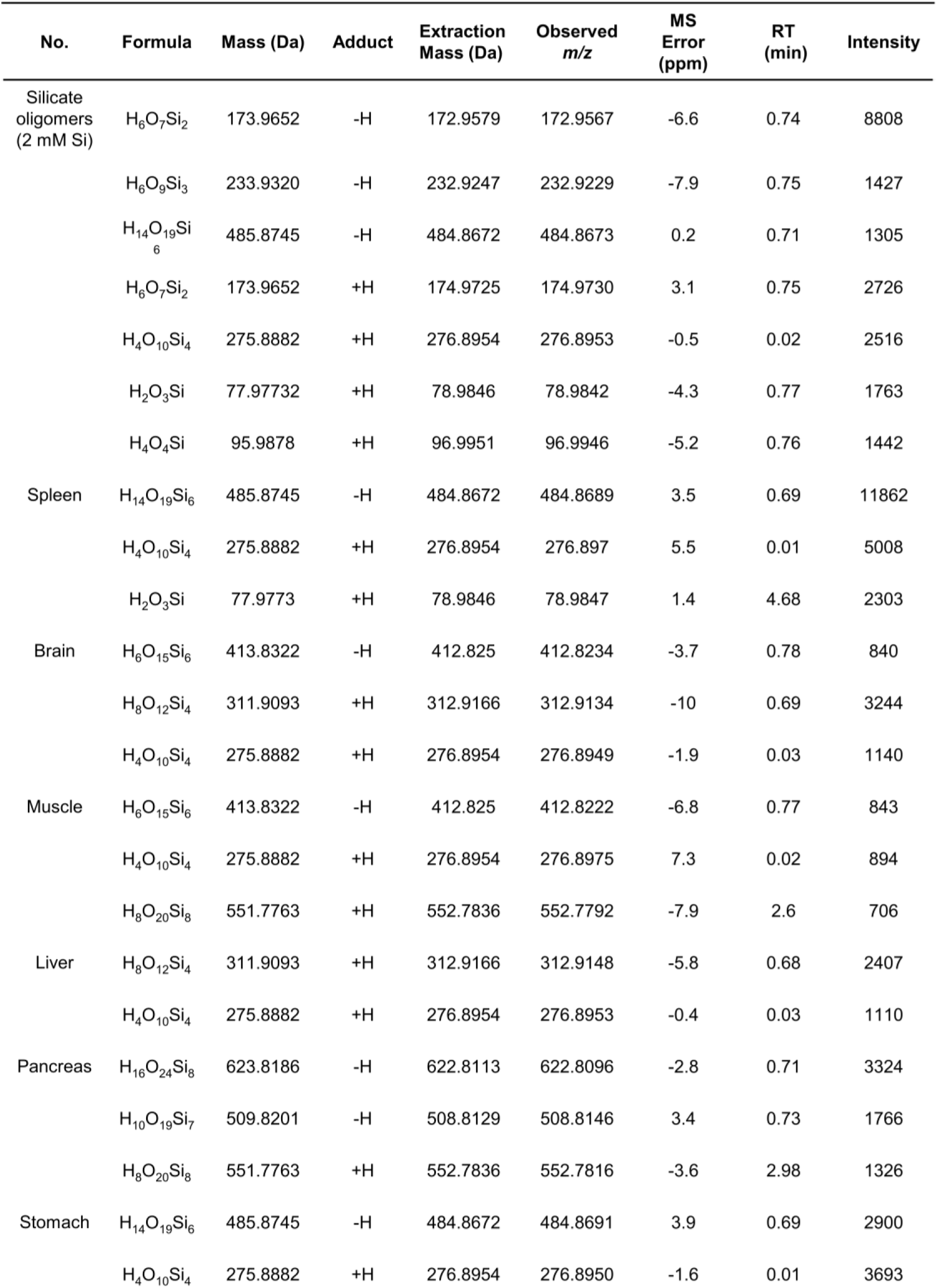
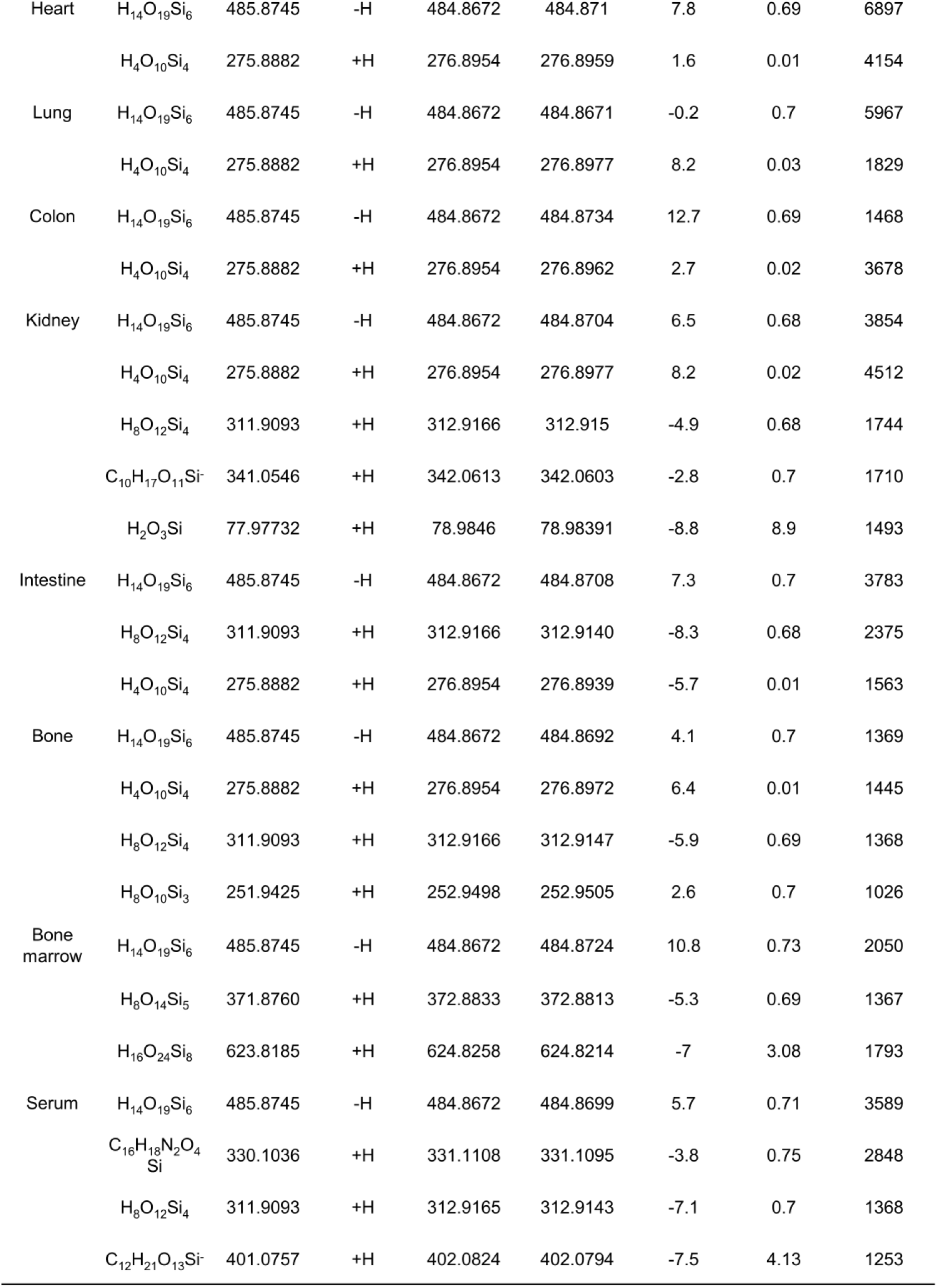
Related to Table 2. Potential silicon compounds in mouse tissues and serum detected by UPLC/Triple-TOF-MS.

| No. | Formula | Mass (Da) | Adduct | Extraction Mass (Da) | Observed *m/z* | MS Error (ppm) | RT (min) | Intensity |
| --- | --- | --- | --- | --- | --- | --- | --- | --- |
| Silicate oligomers (2 mM Si) | H<sub>6</sub>O<sub>7</sub>Si<sub>2</sub> | 173.9652 | -H | 172.9579 | 172.9567 | -6.6 | 0.74 | 8808 |
|  | H<sub>6</sub>O<sub>9</sub>Si<sub>3</sub> | 233.9320 | -H | 232.9247 | 232.9229 | -7.9 | 0.75 | 1427 |
|  | H<sub>14</sub>O<sub>19</sub>Si<sub>6</sub> | 485.8745 | -H | 484.8672 | 484.8673 | 0.2 | 0.71 | 1305 |
|  | H<sub>6</sub>O<sub>7</sub>Si<sub>2</sub> | 173.9652 | +H | 174.9725 | 174.9730 | 3.1 | 0.75 | 2726 |
|  | H<sub>4</sub>O<sub>10</sub>Si<sub>4</sub> | 275.8882 | +H | 276.8954 | 276.8953 | -0.5 | 0.02 | 2516 |
|  | H<sub>2</sub>O<sub>3</sub>Si | 77.97732 | +H | 78.9846 | 78.9842 | -4.3 | 0.77 | 1763 |
|  | H<sub>4</sub>O<sub>4</sub>Si | 95.9878 | +H | 96.9951 | 96.9946 | -5.2 | 0.76 | 1442 |
| Spleen | H<sub>14</sub>O<sub>19</sub>Si<sub>6</sub> | 485.8745 | -H | 484.8672 | 484.8689 | 3.5 | 0.69 | 11862 |
|  | H<sub>4</sub>O<sub>10</sub>Si<sub>4</sub> | 275.8882 | +H | 276.8954 | 276.897 | 5.5 | 0.01 | 5008 |
|  | H<sub>2</sub>O<sub>3</sub>Si | 77.9773 | +H | 78.9846 | 78.9847 | 1.4 | 4.68 | 2303 |
| Brain | H<sub>6</sub>O<sub>15</sub>Si<sub>6</sub> | 413.8322 | -H | 412.825 | 412.8234 | -3.7 | 0.78 | 840 |
|  | H<sub>8</sub>O<sub>12</sub>Si<sub>4</sub> | 311.9093 | +H | 312.9166 | 312.9134 | -10 | 0.69 | 3244 |
|  | H<sub>4</sub>O<sub>10</sub>Si<sub>4</sub> | 275.8882 | +H | 276.8954 | 276.8949 | -1.9 | 0.03 | 1140 |
| Muscle | H<sub>6</sub>O<sub>15</sub>Si<sub>6</sub> | 413.8322 | -H | 412.825 | 412.8222 | -6.8 | 0.77 | 843 |
|  | H<sub>4</sub>O<sub>10</sub>Si<sub>4</sub> | 275.8882 | +H | 276.8954 | 276.8975 | 7.3 | 0.02 | 894 |
|  | H<sub>8</sub>O<sub>20</sub>Si<sub>8</sub> | 551.7763 | +H | 552.7836 | 552.7792 | -7.9 | 2.6 | 706 |
| Liver | H<sub>8</sub>O<sub>12</sub>Si<sub>4</sub> | 311.9093 | +H | 312.9166 | 312.9148 | -5.8 | 0.68 | 2407 |
|  | H<sub>4</sub>O<sub>10</sub>Si<sub>4</sub> | 275.8882 | +H | 276.8954 | 276.8953 | -0.4 | 0.03 | 1110 |
| Pancreas | H<sub>16</sub>O<sub>24</sub>Si<sub>8</sub> | 623.8186 | -H | 622.8113 | 622.8096 | -2.8 | 0.71 | 3324 |
|  | H<sub>10</sub>O<sub>19</sub>Si<sub>7</sub> | 509.8201 | -H | 508.8129 | 508.8146 | 3.4 | 0.73 | 1766 |
|  | H<sub>8</sub>O<sub>20</sub>Si<sub>8</sub> | 551.7763 | +H | 552.7836 | 552.7816 | -3.6 | 2.98 | 1326 |
| Stomach | H<sub>14</sub>O<sub>19</sub>Si<sub>6</sub> | 485.8745 | -H | 484.8672 | 484.8691 | 3.9 | 0.69 | 2900 |
|  | H<sub>4</sub>O<sub>10</sub>Si<sub>4</sub> | 275.8882 | +H | 276.8954 | 276.8950 | -1.6 | 0.01 | 3693 |
| Heart | H<sub>14</sub>O<sub>19</sub>Si<sub>6</sub> | 485.8745 | -H | 484.8672 | 484.871 | 7.8 | 0.69 | 6897 |
|  | H<sub>4</sub>O<sub>10</sub>Si<sub>4</sub> | 275.8882 | +H | 276.8954 | 276.8959 | 1.6 | 0.01 | 4154 |
| Lung | H<sub>14</sub>O<sub>19</sub>Si<sub>6</sub> | 485.8745 | -H | 484.8672 | 484.8671 | -0.2 | 0.7 | 5967 |
|  | H<sub>4</sub>O<sub>10</sub>Si<sub>4</sub> | 275.8882 | +H | 276.8954 | 276.8977 | 8.2 | 0.03 | 1829 |
| Colon | H<sub>14</sub>O<sub>19</sub>Si<sub>6</sub> | 485.8745 | -H | 484.8672 | 484.8734 | 12.7 | 0.69 | 1468 |
|  | H<sub>4</sub>O<sub>10</sub>Si<sub>4</sub> | 275.8882 | +H | 276.8954 | 276.8962 | 2.7 | 0.02 | 3678 |
| Kidney | H<sub>14</sub>O<sub>19</sub>Si<sub>6</sub> | 485.8745 | -H | 484.8672 | 484.8704 | 6.5 | 0.68 | 3854 |
|  | H<sub>4</sub>O<sub>10</sub>Si<sub>4</sub> | 275.8882 | +H | 276.8954 | 276.8977 | 8.2 | 0.02 | 4512 |
|  | H<sub>8</sub>O<sub>12</sub>Si<sub>4</sub> | 311.9093 | +H | 312.9166 | 312.915 | -4.9 | 0.68 | 1744 |
|  | C<sub>10</sub>H<sub>17</sub>O<sub>11</sub>Si<sup>-</sup> | 341.0546 | +H | 342.0613 | 342.0603 | -2.8 | 0.7 | 1710 |
|  | H<sub>2</sub>O<sub>3</sub>Si | 77.97732 | +H | 78.9846 | 78.98391 | -8.8 | 8.9 | 1493 |
| Intestine | H<sub>14</sub>O<sub>19</sub>Si<sub>6</sub> | 485.8745 | -H | 484.8672 | 484.8708 | 7.3 | 0.7 | 3783 |
|  | H<sub>8</sub>O<sub>12</sub>Si<sub>4</sub> | 311.9093 | +H | 312.9166 | 312.9140 | -8.3 | 0.68 | 2375 |
|  | H<sub>4</sub>O<sub>10</sub>Si<sub>4</sub> | 275.8882 | +H | 276.8954 | 276.8939 | -5.7 | 0.01 | 1563 |
| Bone | H<sub>14</sub>O<sub>19</sub>Si<sub>6</sub> | 485.8745 | -H | 484.8672 | 484.8692 | 4.1 | 0.7 | 1369 |
|  | H<sub>4</sub>O<sub>10</sub>Si<sub>4</sub> | 275.8882 | +H | 276.8954 | 276.8972 | 6.4 | 0.01 | 1445 |
|  | H<sub>8</sub>O<sub>12</sub>Si<sub>4</sub> | 311.9093 | +H | 312.9166 | 312.9147 | -5.9 | 0.69 | 1368 |
|  | H<sub>8</sub>O<sub>10</sub>Si<sub>3</sub> | 251.9425 | +H | 252.9498 | 252.9505 | 2.6 | 0.7 | 1026 |
| Bone marrow | H<sub>14</sub>O<sub>19</sub>Si<sub>6</sub> | 485.8745 | -H | 484.8672 | 484.8724 | 10.8 | 0.73 | 2050 |
|  | H<sub>8</sub>O<sub>14</sub>Si<sub>5</sub> | 371.8760 | +H | 372.8833 | 372.8813 | -5.3 | 0.69 | 1367 |
|  | H<sub>16</sub>O<sub>24</sub>Si<sub>8</sub> | 623.8185 | +H | 624.8258 | 624.8214 | -7 | 3.08 | 1793 |
| Serum | H<sub>14</sub>O<sub>19</sub>Si<sub>6</sub> | 485.8745 | -H | 484.8672 | 484.8699 | 5.7 | 0.71 | 3589 |
|  | C<sub>16</sub>H<sub>18</sub>N<sub>2</sub>O<sub>4</sub>Si | 330.1036 | +H | 331.1108 | 331.1095 | -3.8 | 0.75 | 2848 |
|  | H<sub>8</sub>O<sub>12</sub>Si<sub>4</sub> | 311.9093 | +H | 312.9165 | 312.9143 | -7.1 | 0.7 | 1368 |
|  | C<sub>12</sub>H<sub>21</sub>O<sub>13</sub>Si<sup>-</sup> | 401.0757 | +H | 402.0824 | 402.0794 | -7.5 | 4.13 | 1253 |

**Table S4.**
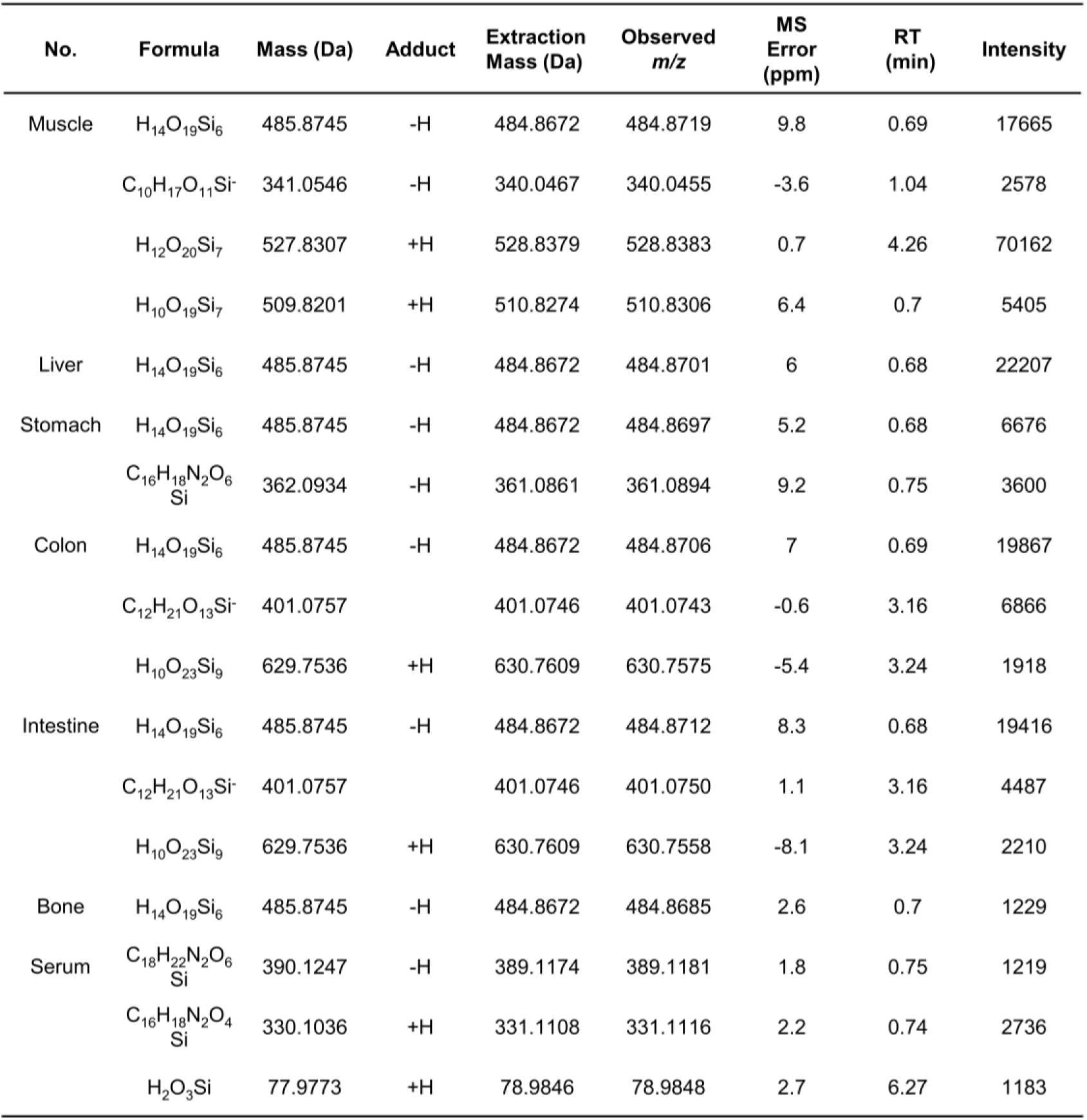
Related to Table 3. Potential silicon compounds in human tissues and serum detected by UPLC/Triple-TOF-MS.

