## Supplementary information for "Quantitative Prevalence, Chemical Speciation, and ECM-Embedded Networks of Biogenic Silicon in Animal Tissues Revealed by Refined Analytical Methods"

Statistical analysis was performed with SPSS (version 25.0) or GraphPad Prism (version 8.0) software. All values in graphs are presented as mean±SEM.

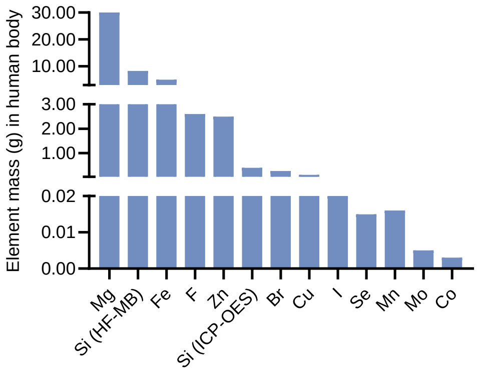


**Figure S1**. Related to Figure 1. Element mass in the human body (70 kg). Elements that constitute more than 0.01% of the human body are known as major elements, including C, H, O, N, K, S, Na, Cl, Mg, Ca, and P. Those that make up less than 0.01% are referred to as trace elements, including Si, Fe, F, Zn, Br, Cu, I, Se, Mn, Mo, Co. The trace element mass values are sourced from literature reports, whereas the silicon mass value is based on measurements conducted in this study.

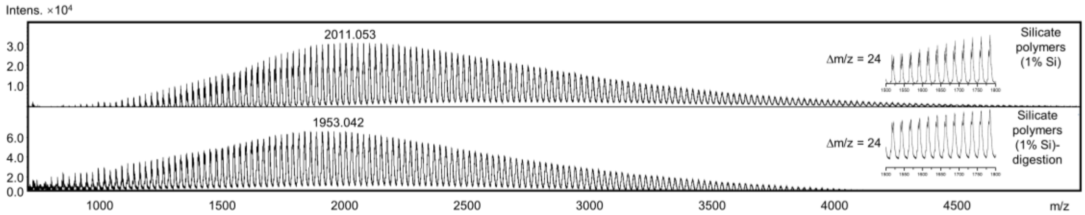


**Figure S2**. Related to Figure 2. Potential silicate polymers in synthetic silicate polymers or after enzyme digestion identified by MALDI-TOF-MS.

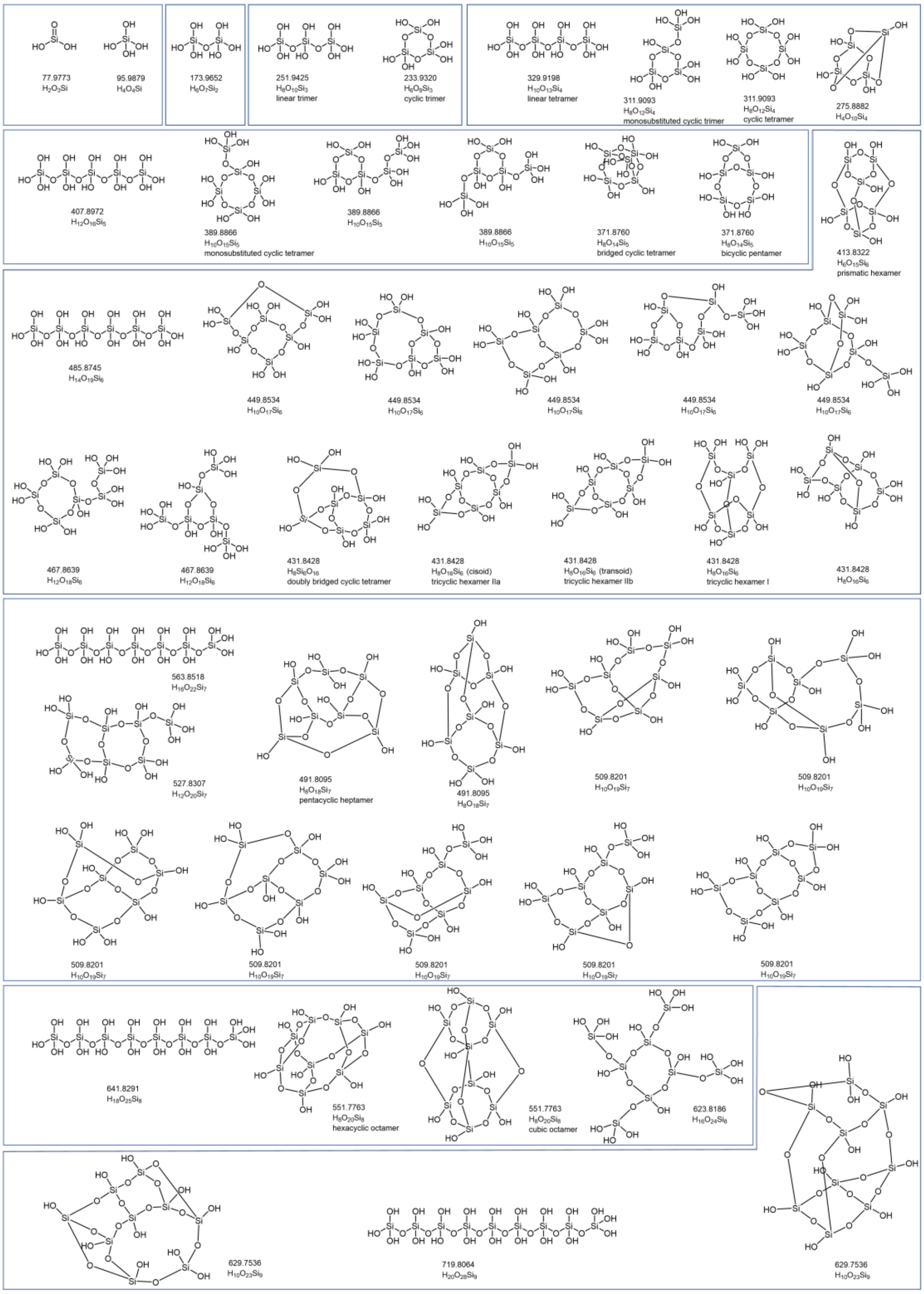

**Figure S3**. Related to Figure 2. Chemical structural formula library of silicate oligomers.

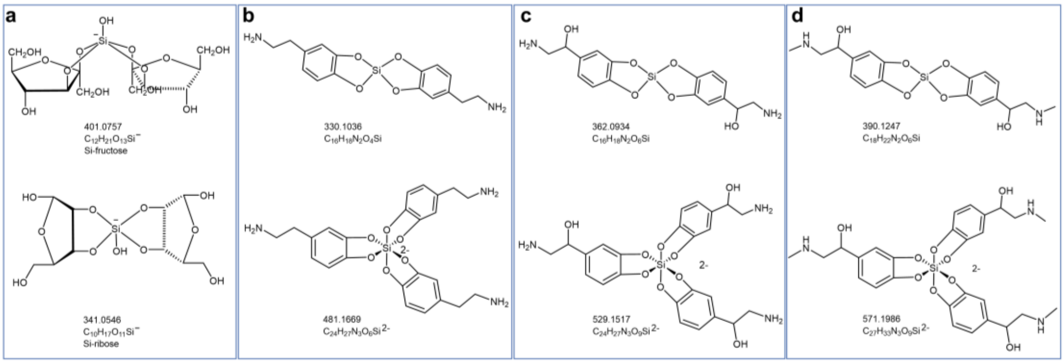


**Figure S4**. Related to Figure 2. Chemical structural formula library of sugar silicates (**a**), dopamine-silicates (**b**), norepinephrine-silicates (**c**), and epinephrine-silicates (**d**).

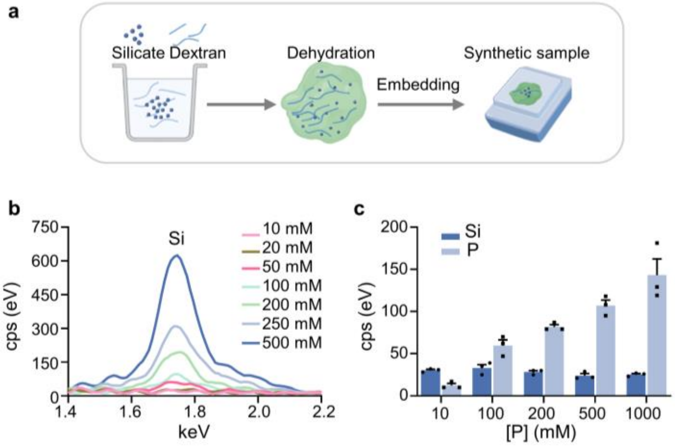


**Figure S5**. Related to Figure 3. **a**, Schematic of the micro-XRF experimental setup for silicon detection in synthetic samples. 100 mg of dextran powder is dissolved in 1 mL of sodium silicate solution, then lyophilized, embedded in paraffin and cut the surface flat as the synthetic samples. **b**, Silicon peaks in XRF spectra of different concentrations of sodium silicate encapsulated in synthetic samples. **c**, Silicon peak intensity in XRF spectra of different concentrations of sodium phosphate encapsulated in synthetic samples containing 20 mM sodium silicate.

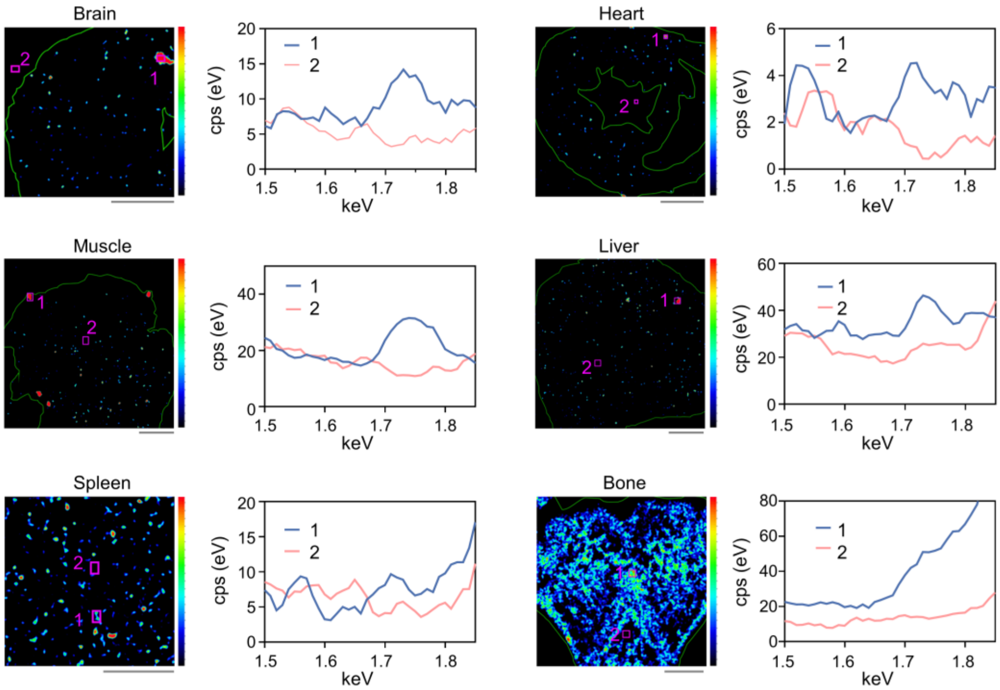


**Figure S6**. Related to Figure 3. Micro-XRF spectra (right) showing the silicon peaks, taken from the areas indicated by pink boxes in the micro-XRF heat maps (left) of different mouse tissues (brain, heart, muscle, liver, spleen, and rat bone). Scale bars, 1000 µm.

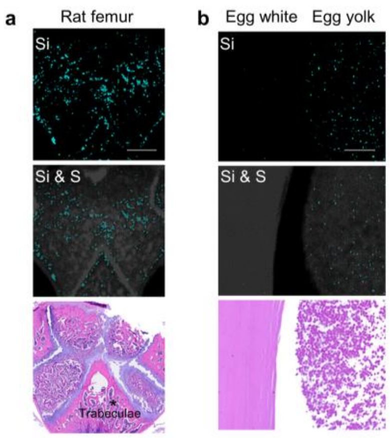


**Figure S7**. Related to Figure 3. Micro-XRF images are collected from rat femur, egg white and yolk. Scale bars, 1000 μm.

**Table S1.** Related to Figure 1. Quantification of silicon in food determined by HF-MB method (n = 3).

| Food | MB (μg/g, wet weight) | MB (μg/g, dry weight) | Food | MB (μg/g, wet weight) | MB (μg/g, dry weight) |
|---|---|---|---|---|---|
| **Beef** | 165.3±37.2 | 924.0±134.3 | **Onion** | 35.6±3.9 | 471.0±125.5 |
| **Chicken** | 150.3±9.6 | 698.1±43.6 | **Cabbage** | 46.5±3.2 | 580.5±56.0 |
| **Lamb** | 195.9±24.6 | 975.8±21.7 | **Lettuce** | 52.9±8.4 | 897.8±55.9 |
| **Pork** | 149.1±13.9 | 800.3±164.9 | **Okra** | 49.8±3.9 | 488.8±34.9 |
| **Fish** | 102.5±14.7 | 694.4±76.6 | **Potato** | 25.1±1.2 | 143.7±15.6 |
| **Tomato** | 14.7±1.1 | 410.8±127.1 | **Garlic** | 30.7±1.8 | 82.9±5.6 |

**Table S2.** Related to Table 2. Chemical structure of potential silicon compounds in synthetic silicate oligomers solution (2 mM sodium silicate solution) identified by UPLC/Triple-TOF-MS.

| No. | Chemical Structure of silicon compound | | | | | |
|---|---|---|---|---|---|---|
| Silicate oligomers (2 mM Si) | 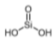<br>77.9773<br>$H_2O_3Si$ | 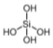<br>95.9879<br>$H_4O_4Si$ | 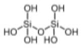<br>173.9652<br>$H_6O_7Si_2$ | 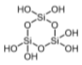<br>233.9320<br>$H_6O_9Si_3$ | 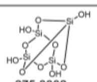<br>275.8882<br>$H_4O_{10}Si_4$ | 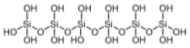<br>485.8745<br>$H_{14}O_{19}Si_6$ |

**Table S3**. Related to Table 2. Potential silicon compounds in mouse tissues and serum detected by UPLC/Triple-TOF-MS.

| No. | Formula | Mass (Da) | Adduct | Extraction Mass (Da) | Observed *m/z* | MS Error (ppm) | RT (min) | Intensity |
|---|---|---|---|---|---|---|---|---|
| Silicate oligomers (2 mM Si) | $H_6O_7Si_2$ | 173.9652 | -H | 172.9579 | 172.9567 | -6.6 | 0.74 | 8808 |
| | $H_6O_9Si_3$ | 233.9320 | -H | 232.9247 | 232.9229 | -7.9 | 0.75 | 1427 |
| | $H_{14}O_{19}Si_6$ | 485.8745 | -H | 484.8672 | 484.8673 | 0.2 | 0.71 | 1305 |
| | $H_6O_7Si_2$ | 173.9652 | +H | 174.9725 | 174.9730 | 3.1 | 0.75 | 2726 |
| | $H_4O_{10}Si_4$ | 275.8882 | +H | 276.8954 | 276.8953 | -0.5 | 0.02 | 2516 |
| | $H_2O_3Si$ | 77.97732 | +H | 78.9846 | 78.9842 | -4.3 | 0.77 | 1763 |
| | $H_4O_4Si$ | 95.9878 | +H | 96.9951 | 96.9946 | -5.2 | 0.76 | 1442 |
| Spleen | $H_{14}O_{19}Si_6$ | 485.8745 | -H | 484.8672 | 484.8689 | 3.5 | 0.69 | 11862 |
| | $H_4O_{10}Si_4$ | 275.8882 | +H | 276.8954 | 276.897 | 5.5 | 0.01 | 5008 |
| | $H_2O_3Si$ | 77.9773 | +H | 78.9846 | 78.9847 | 1.4 | 4.68 | 2303 |
| Brain | $H_6O_{15}Si_6$ | 413.8322 | -H | 412.825 | 412.8234 | -3.7 | 0.78 | 840 |
| | $H_8O_{12}Si_4$ | 311.9093 | +H | 312.9166 | 312.9134 | -10 | 0.69 | 3244 |
| | $H_4O_{10}Si_4$ | 275.8882 | +H | 276.8954 | 276.8949 | -1.9 | 0.03 | 1140 |
| Muscle | $H_6O_{15}Si_6$ | 413.8322 | -H | 412.825 | 412.8222 | -6.8 | 0.77 | 843 |
| | $H_4O_{10}Si_4$ | 275.8882 | +H | 276.8954 | 276.8975 | 7.3 | 0.02 | 894 |
| | $H_8O_{20}Si_8$ | 551.7763 | +H | 552.7836 | 552.7792 | -7.9 | 2.6 | 706 |
| Liver | $H_8O_{12}Si_4$ | 311.9093 | +H | 312.9166 | 312.9148 | -5.8 | 0.68 | 2407 |
| | $H_4O_{10}Si_4$ | 275.8882 | +H | 276.8954 | 276.8953 | -0.4 | 0.03 | 1110 |
| Pancreas | $H_{16}O_{24}Si_8$ | 623.8186 | -H | 622.8113 | 622.8096 | -2.8 | 0.71 | 3324 |
| | $H_{10}O_{19}Si_7$ | 509.8201 | -H | 508.8129 | 508.8146 | 3.4 | 0.73 | 1766 |
| | $H_8O_{20}Si_8$ | 551.7763 | +H | 552.7836 | 552.7816 | -3.6 | 2.98 | 1326 |
| Stomach | $H_{14}O_{19}Si_6$ | 485.8745 | -H | 484.8672 | 484.8691 | 3.9 | 0.69 | 2900 |
| | $H_4O_{10}Si_4$ | 275.8882 | +H | 276.8954 | 276.8950 | -1.6 | 0.01 | 3693 |

| | | | | | | | | |
|---|---|---|---|---|---|---|---|---|
| Heart | $H_{14}O_{19}Si_6$ | 485.8745 | -H | 484.8672 | 484.871 | 7.8 | 0.69 | 6897 |
| | $H_4O_{10}Si_4$ | 275.8882 | +H | 276.8954 | 276.8959 | 1.6 | 0.01 | 4154 |
| Lung | $H_{14}O_{19}Si_6$ | 485.8745 | -H | 484.8672 | 484.8671 | -0.2 | 0.7 | 5967 |
| | $H_4O_{10}Si_4$ | 275.8882 | +H | 276.8954 | 276.8977 | 8.2 | 0.03 | 1829 |
| Colon | $H_{14}O_{19}Si_6$ | 485.8745 | -H | 484.8672 | 484.8734 | 12.7 | 0.69 | 1468 |
| | $H_4O_{10}Si_4$ | 275.8882 | +H | 276.8954 | 276.8962 | 2.7 | 0.02 | 3678 |
| Kidney | $H_{14}O_{19}Si_6$ | 485.8745 | -H | 484.8672 | 484.8704 | 6.5 | 0.68 | 3854 |
| | $H_4O_{10}Si_4$ | 275.8882 | +H | 276.8954 | 276.8977 | 8.2 | 0.02 | 4512 |
| | $H_8O_{12}Si_4$ | 311.9093 | +H | 312.9166 | 312.915 | -4.9 | 0.68 | 1744 |
| | $C_{10}H_{17}O_{11}Si^-$ | 341.0546 | +H | 342.0613 | 342.0603 | -2.8 | 0.7 | 1710 |
| | $H_2O_3Si$ | 77.97732 | +H | 78.9846 | 78.98391 | -8.8 | 8.9 | 1493 |
| Intestine | $H_{14}O_{19}Si_6$ | 485.8745 | -H | 484.8672 | 484.8708 | 7.3 | 0.7 | 3783 |
| | $H_8O_{12}Si_4$ | 311.9093 | +H | 312.9166 | 312.9140 | -8.3 | 0.68 | 2375 |
| | $H_4O_{10}Si_4$ | 275.8882 | +H | 276.8954 | 276.8939 | -5.7 | 0.01 | 1563 |
| Bone | $H_{14}O_{19}Si_6$ | 485.8745 | -H | 484.8672 | 484.8692 | 4.1 | 0.7 | 1369 |
| | $H_4O_{10}Si_4$ | 275.8882 | +H | 276.8954 | 276.8972 | 6.4 | 0.01 | 1445 |
| | $H_8O_{12}Si_4$ | 311.9093 | +H | 312.9166 | 312.9147 | -5.9 | 0.69 | 1368 |
| | $H_8O_{10}Si_3$ | 251.9425 | +H | 252.9498 | 252.9505 | 2.6 | 0.7 | 1026 |
| Bone marrow | $H_{14}O_{19}Si_6$ | 485.8745 | -H | 484.8672 | 484.8724 | 10.8 | 0.73 | 2050 |
| | $H_8O_{14}Si_5$ | 371.8760 | +H | 372.8833 | 372.8813 | -5.3 | 0.69 | 1367 |
| | $H_{16}O_{24}Si_8$ | 623.8185 | +H | 624.8258 | 624.8214 | -7 | 3.08 | 1793 |
| Serum | $H_{14}O_{19}Si_6$ | 485.8745 | -H | 484.8672 | 484.8699 | 5.7 | 0.71 | 3589 |
| | $C_{16}H_{18}N_2O_4Si$ | 330.1036 | +H | 331.1108 | 331.1095 | -3.8 | 0.75 | 2848 |
| | $H_8O_{12}Si_4$ | 311.9093 | +H | 312.9165 | 312.9143 | -7.1 | 0.7 | 1368 |
| | $C_{12}H_{21}O_{13}Si^-$ | 401.0757 | +H | 402.0824 | 402.0794 | -7.5 | 4.13 | 1253 |

**Table S4**. Related to Table 3. Potential silicon compounds in human tissues and serum detected by UPLC/Triple-TOF-MS.

| No. | Formula | Mass (Da) | Adduct | Extraction Mass (Da) | Observed *m/z* | MS Error (ppm) | RT (min) | Intensity |
|---|---|---|---|---|---|---|---|---|
| Muscle | $H_{14}O_{19}Si_6$ | 485.8745 | -H | 484.8672 | 484.8719 | 9.8 | 0.69 | 17665 |
| | $C_{10}H_{17}O_{11}Si^-$ | 341.0546 | -H | 340.0467 | 340.0455 | -3.6 | 1.04 | 2578 |
| | $H_{12}O_{20}Si_7$ | 527.8307 | +H | 528.8379 | 528.8383 | 0.7 | 4.26 | 70162 |
| | $H_{10}O_{19}Si_7$ | 509.8201 | +H | 510.8274 | 510.8306 | 6.4 | 0.7 | 5405 |
| Liver | $H_{14}O_{19}Si_6$ | 485.8745 | -H | 484.8672 | 484.8701 | 6 | 0.68 | 22207 |
| Stomach | $H_{14}O_{19}Si_6$ | 485.8745 | -H | 484.8672 | 484.8697 | 5.2 | 0.68 | 6676 |
| | $C_{16}H_{18}N_2O_6Si$ | 362.0934 | -H | 361.0861 | 361.0894 | 9.2 | 0.75 | 3600 |
| Colon | $H_{14}O_{19}Si_6$ | 485.8745 | -H | 484.8672 | 484.8706 | 7 | 0.69 | 19867 |
| | $C_{12}H_{21}O_{13}Si^-$ | 401.0757 | | 401.0746 | 401.0743 | -0.6 | 3.16 | 6866 |
| | $H_{10}O_{23}Si_9$ | 629.7536 | +H | 630.7609 | 630.7575 | -5.4 | 3.24 | 1918 |
| Intestine | $H_{14}O_{19}Si_6$ | 485.8745 | -H | 484.8672 | 484.8712 | 8.3 | 0.68 | 19416 |
| | $C_{12}H_{21}O_{13}Si^-$ | 401.0757 | | 401.0746 | 401.0750 | 1.1 | 3.16 | 4487 |
| | $H_{10}O_{23}Si_9$ | 629.7536 | +H | 630.7609 | 630.7558 | -8.1 | 3.24 | 2210 |
| Bone | $H_{14}O_{19}Si_6$ | 485.8745 | -H | 484.8672 | 484.8685 | 2.6 | 0.7 | 1229 |
| Serum | $C_{18}H_{22}N_2O_6Si$ | 390.1247 | -H | 389.1174 | 389.1181 | 1.8 | 0.75 | 1219 |
| | $C_{16}H_{18}N_2O_4Si$ | 330.1036 | +H | 331.1108 | 331.1116 | 2.2 | 0.74 | 2736 |
| | $H_2O_3Si$ | 77.9773 | +H | 78.9846 | 78.9848 | 2.7 | 6.27 | 1183 |
